# Molecular insights into the gating mechanisms of voltage-gated calcium channel Ca_V_2.3

**DOI:** 10.1101/2022.12.19.521133

**Authors:** Yiwei Gao, Shuai Xu, Xiaoli Cui, Hao Xu, Yunlong Qiu, Yiqing Wei, Yanli Dong, Boling Zhu, Chao Peng, Shiqi Liu, Xuejun Cai Zhang, Jianyuan Sun, Zhuo Huang, Yan Zhao

## Abstract

High-voltage-activated R-type Ca_V_2.3 channel plays pivotal roles in many physiological activities and is implicated in epilepsy, convulsions, and other neurodevelopmental impairments. Here, we determine the high-resolution cryo-electron microscopy (cryo-EM) structure of human Ca_V_2.3 in complex with the α2δ1 and β1 subunits. The VSD_II_ is stabilized in the resting state. Electrophysiological experiments elucidate that the conformational change of VSD_II_ in response to variation in membrane potential is not required for channel activation, whereas the other VSDs are essential for channel opening. The intracellular gate is blocked by the W-helix. A pre-W-helix adjacent to the W-helix can significantly regulate closed-state inactivation (CSI) by modulating the association and dissociation of the W-helix with the gate. Electrostatic interactions formed between the negatively charged domain on S6_II_, which is exclusively conserved in the Ca_V_2 family, and nearby regions at the alpha-interacting domain (AID) and S4-S5_II_ helix are identified. Further functional analyses indicate that these interactions are critical for the open-state inactivation (OSI) of Ca_V_2 channels.

## Introduction

Voltage-gated calcium (Ca_V_) channels mediate calcium influx to cells in response to changes in membrane potential^1-3^. Their cellular roles have been emphasized for decades in a variety of studies and include hormone secretion^4,5^, neurotransmitter release^6,7^, and muscle contraction^8,9^. Ca_V_ channel members are categorized into the Ca_V_1, Ca_V_2, and Ca_V_3 subfamilies based on sequence identity or alternatively classified into T-, L-, P/Q-, N-, and R-types according to their pharmacological and biophysical profiles^1^. The so-called pharmacoresistant (R-type) Ca_V_2.3 is widely expressed in the brain and enriched in the hippocampus, cerebral cortex, amygdala, and corpus striatum^10-12^. Electrophysiological investigations revealed that currents mediated by Ca_V_2.3 are resistant to common Ca_V_ blockers or gating modifiers such as nifedipine, nimodipine, ω-Aga-IVA, etc^13^. Ca_V_2.3 channels exhibit cumulative inactivation in response to brief and repetitive depolarizations, a process known as preferential closed-state inactivation (CSI)^14^. Furthermore, Ca_V_2.3 is involved in a broad spectrum of neuronal activities^10,12,15^. Previous studies have reported that Ca_V_2.3 participates in multiple physiological processes in the central nervous system, such as inducing long-term potentiation (LTP) and post-tetanic potentiation in mossy fiber synapses^16^, modulating the burst firing mode of action potentials^17,18^, and regulating synaptic strength in hippocampal CA1 pyramidal neurons^19^. In recent years, increasing evidence has revealed that dysfunction of Ca_V_2.3 is linked to epilepsy^20,21^, convulsions^22,23^, and neurodevelopmental impairments^24^, suggesting that Ca_V_2.3 is a pivotal player in the pathogenesis of a series of neurological disorders.

The molecular basis of Ca_V_ channels has been investigated extensively over the past several decades, including structural studies of L-type Ca_V_1.1^25,26^, N-type Ca_V_2.2^27,28^, and T-type Ca_V_3.1^29^ and Ca_V_3.3^30^ in the apo form or distinct modulator-bound states. These structures provide substantial insights into the architecture, subunit assembly, and modulator actions of the Ca_V_ channels. However, the gating mechanism of Ca_V_ channels is still far from fully understood. For instance, in the Ca_V_2.2 structure, the VSD_II_ is trapped in a resting state by a PIP_2_ molecule at a membrane potential of ∼0 mV^27,28^. The functional roles of the VSD_II_ trapped in the resting state by PIP_2_ remain unknown. A considerable number of pathogenic mutations have been identified in the VSDs of neuronal Ca_V_2 channels, demonstrating that VSD dysfunctions contribute to the genesis of spinocerebellar ataxia (SCA), episodic ataxia (EA), and familial hemiplegic migraine (FHM)^31^. Moreover, the Ca_V_2.2 and Ca_V_2.3 channels inactivate preferentially from the intermediate closed state along the activation pathway, which is important in controlling the short-term dynamics of synaptic efficacy^14,32^. In our previous study, we elucidated that residue W768 on the W-helix located within the DII-III linker serves as a key determinant of the CSI of the Ca_V_2.2 channel^28^. However, Ca_V_2.3 is characterized by a more prominent preferential CSI than Ca_V_2.2^14^. It is also interesting to explore the modulation mechanism of CSI in Ca_V_2.3. Furthermore, the high-voltage-activated (HVA) Ca_V_1 and Ca_V_2 channels harbor a conserved α-helix connecting Domain I and Domain II (alpha-interaction domain, or AID). Previous studies have indicated that AID might contribute to the open-state inactivation (OSI) of the HVA Ca_V_ channels^33,34^. However, the inactivation properties of Ca_V_1 and Ca_V_2 channels are dramatically different^33^. Mechanistic insight into the inactivation processes of the HVA Ca_V_ channels will help us to fully uncover the physiological role of Ca_V_ channels and facilitate the development of therapeutic solutions for Ca_V_-related diseases.

In this study, we expressed and purified human Ca_V_2.3 in complex with auxiliary subunits α2δ1 and β1 and unveiled the high-resolution structure of this protein complex. Further mutagenesis and electrophysiological experiments were performed. Our results provide insights into the pharmacological resistance properties of Ca_V_2.3, the asynchronous functional roles of the VSDs, the mechanism by which the pre-W-helix regulates the CSI, and the OSI process modulated by the negatively charged domain on S6_II_ (S6_II_^NCD^).

## Results and Discussion

### Architecture of the Ca_V_2.3 complex

To gain structural insights into the Ca_V_2.3 complex, we expressed and purified full-length wild-type human Ca_V_2.3 α1E subunit (CACNA1E), α2δ1 (CACNA2D1) and β1 (CACB1) using a HEK 293-F expression system. The Ca_V_2.3-α2δ1-β1 complex was solubilized using n-Dodecyl-βD-maltoside (DDM) and purified using a strep-tactin affinity column, followed by further purification by size-exclusion chromatography (SEC) in a running buffer containing glycol-diosgenin (GDN) to remove protein aggregates (Supplementary Figure 1a, see Method section for details). The peak fractions were subsequently collected and concentrated for cryo-EM sample preparation (Supplementary Figure 1b). A total of 2,096 micrographs were collected. Data processing of the dataset gave rise to a 3.1-Å cryo-EM map, which is rich in high-resolution structural features, including densities for side chains, lipid molecules and glycosylations (Figure 1, Supplementary Figure 2, and Table S1).

**Figure 1.**
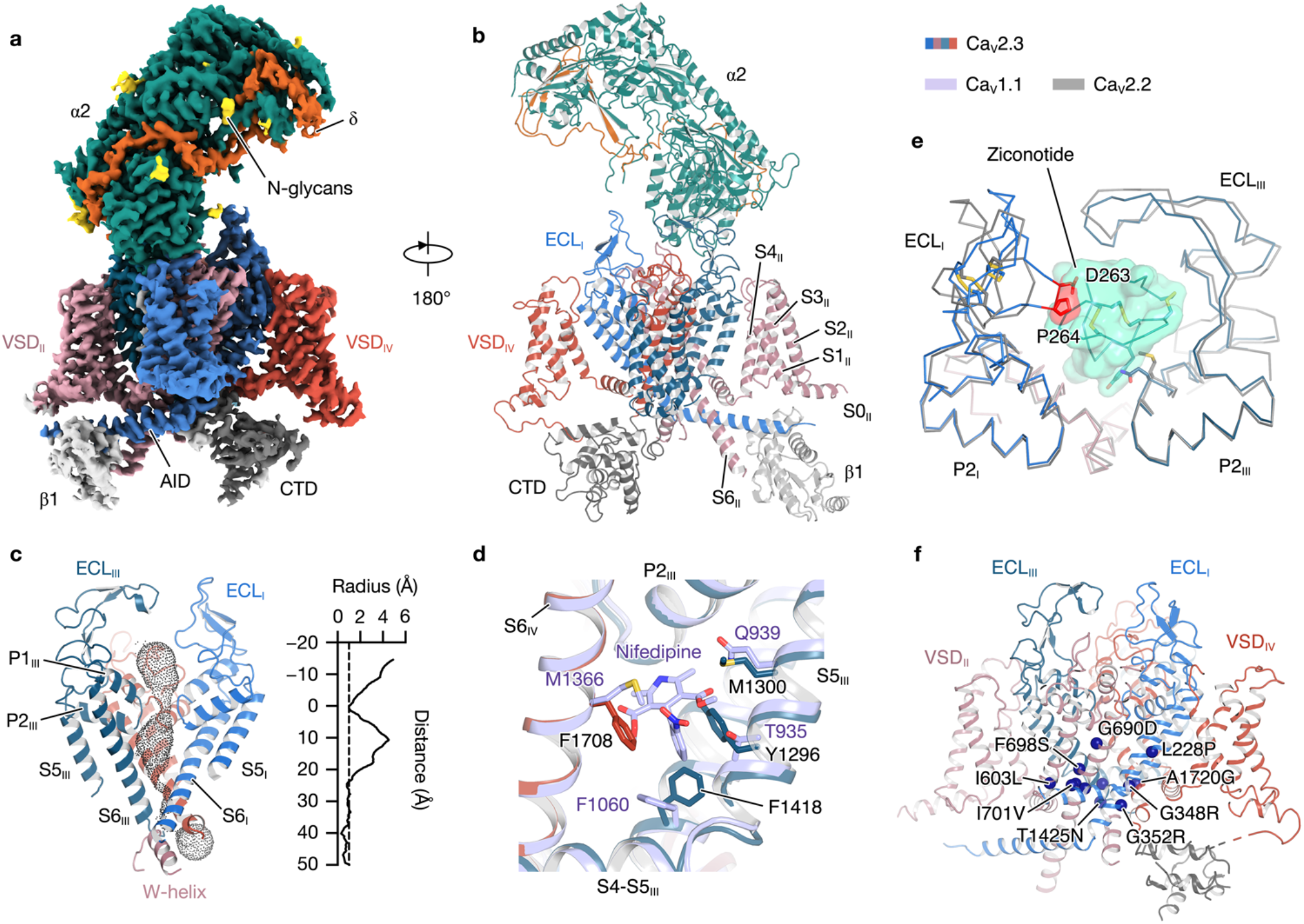
Architecture of the Ca_V_2.3-α2δ1-β1 complex. **a**–**b**. Cryo-EM map (**a**) and model (**b**) of the Ca_V_2.3-α2δ1-β1 complex. The Ca_V_2.3 α1E pore-forming subunit are colored in pink, red, blue, and deep cyan. The C-terminal domain (CTD) of Ca_V_2.3 are colored in gray. The auxiliary subunits α2, δ, and β1 are colored in green, orange, and white, respectively. **c**. The ion conducting pathway and pore profiling of the Ca_V_2.3. The ion conducting pathway is viewed in parallel to the membrane and shown in black dots. The radius of the pore is calculated using the HOLE program. The vertical dashed line marks the 1.0-Å pore radius, which represents the ionic radius of calcium. **d**. Superimposition of the DIII-IV fenestration between Ca_V_2.3 and Ca_V_1.1 complex. The nifedipine molecule bound to the Ca_V_1.1 complex is shown as sticks. Residues stabilizing the nifedipine molecule in Ca_V_1.1, and residues which might form steric clashes in Ca_V_2.3 are shown as sticks. **e**. Superimposition between the ‘P-loops’ (P1 and P2) and extracellular loops (ECLs) of the Ca_V_2.3 and the ziconotide-bound Ca_V_2.2. The Ca_V_2.3 and Ca_V_2.2 are shown as ribbon, overlaid with the ziconotide shown as transparent green surfaces. Potential steric clash between the ECL_I_ of Ca_V_2.3 and the ziconotide is highlighted in red. **f**. Pathogenic mutations of the Ca_V_2.3, shown in blue spheres. The four domains of Ca_V_2.3 are colored in pink, red, blue, and deep cyan, respectively. Ca_V_1.1 is colored in purple and Ca_V_2.2 is colored in gray.

The Ca_V_2.3 complex exhibited a conventional shape that resembles that of the Ca_V_2.2 and Ca_V_1.1 complexes (Figure 1). The complex is composed of the transmembrane α1E subunit, extracellular α2δ1 subunit, and intracellular β1 subunit (Figure 1a–1b). The α1E subunit is a pseudotetrameric pore-forming subunit and can be divided into Domain I (DI) to IV (DIV). Each domain of the α1E subunit is composed of 6 transmembrane helices (S1–S6), comprising the voltage-sensing domain (S1–S4) and the pore domain (S5–S6). The P1 and P2 helices located between the S5 and S6 helices formed the selectivity filter (Figure 1c). Similar to other Ca_V_ channels, Ca_V_2.3 harbors four extracellular loops (ECLs) that is also positioned between S5 and S6 helices in the pore domain (Figure 1a–1c). The ECL_I_ and ECL_II_ are critical for the association between the α1 subunit and the α2δ1 subunit (Figure 1a–1b). The S6 helices from the four domains converge on the cytoplasmic side and form the intracellular gate of the channel. In our structure, the intracellular gate was determined in its closed state, in line with the observations from other voltage-gated channels (Figure 1c). Moreover, the closed gate of Ca_V_2.3 is further stabilized by the W-helix from the DII-DIII linker, which is consistent with a previous study on Ca_V_2.2 and indicates that Ca_V_2.3 also adopts the CSI mechanism^28^.

Most Ca_V_ channels serve as pharmaceutical targets of a variety of small-molecule drugs or peptide toxins^13^. However, previous studies indicate that Ca_V_2.3 is resistant to many Ca_V_ modulators, such as nimodipine (L-type), omega-Aga-IVA (P/Q-type), and omega-CTx-GVIA (N-type)^13^. To clarify the structural basis underlying the pharmacoresistance of Ca_V_2.3, we compared the structures of Ca_V_2.3 with Ca_V_1.1 or Ca_V_2.2 in their ligand-bound states (Figure 1d–1e). In the structure of nifedipine-bound Ca_V_1.1, the nifedipine molecule was located within the DIII-DIV fenestration and stabilized by surrounding residues. However, two critical residues are substituted in Ca_V_2.3, namely, Y1296 and F1708. The bulky sidechains of these two residues in Ca_V_2.3 occupy the DIII-DIV fenestration site and thus hinder the binding of nifedipine (Figure 1d). Meanwhile, a previous study reported that the Q1010 of Ca_V_1.2 is important for sensitivity to dihydropyridine (DHP) and the Q1010M mutant had a decreased sensitivity to DHP molecules^35^. The equivalent position in Ca_V_2.3 is occupied by M1300, thus also contributing to the pharmacoresistance of Ca_V_2.3 to DHP molecules. Structural comparison of Ca_V_2.3 and ziconotide-bound Ca_V_2.2 demonstrated that the ECL_I_ loop of Ca_V_2.3 adopts a different conformation, and residues D263 and P264 are placed close to the central axis, giving rise to clashes between the ziconotide and the ECL_I_ of Ca_V_2.3 (Figure 1e). Moreover, other residues on P-loops and ECLs that are involved in ziconotide binding are also not conserved in Ca_V_2.3 (Supplementary Figure 3), rendering Ca_V_2.3 insensitive to the ziconotide.

Gain-of-function mutations and polymorphisms of Ca_V_2.3 channels have already been implicated in the pathological process of developmental and epileptic encephalopathy. Thirteen pathogenic mutations have been identified in Ca_V_2.3^24^. Our high-resolution structure provides a structural template to map all of these pathogenic mutations, which are distributed throughout the complex structure (Figure 1f). Eight of thirteen mutations are located around the intracellular gate, such as I603L, F698S, and I701V, and result in a hyperpolarizing shift in the half-activation voltage^24^ (Figure 1f).

### Functional heterogeneity of the voltage-sensing domains

The voltage-dependent gating characteristics of voltage-gated channels are conferred by their voltage-sensing domains (VSDs). The VSDs of Ca_V_ are conserved helix bundles consisting of S1, S2, S3 and S4 helices (Supplementary Figure 4a). S4 was found to be a positively charged 3_10_ helix, harboring approximately five or six arginines or lysines as gating charges lining one side of the helix at intervals of three residues (Supplementary Figure 4a). The positively charged S4 helices move vertically toward the intracellular or extracellular side of the cell in response to the hyperpolarization or depolarization of the membrane potential. The conformational change of VSD is coupled to the pore domain by a short amphipathic helix S4-S5, which connects the S4 helix of VSD to the S5 helix from the pore domain, thus regulating the transition of the intracellular gate between the open and closed states. Although the four VSDs of Ca_V_ channels are considerably similar in terms of sequence and overall structure, they contribute differentially to the opening of pore^36^.

Superimposition of the structures of Ca_V_2.3 and Ca_V_2.2 revealed that they are comparable overall (r.m.s.d. = 1.46 Å for 2,222 Cα atom pairs). The pore domain was fairly superimposable between Ca_V_2.3 and Ca_V_2.2, including the S5 and S6 helices and extracellular loops (ECLs) I, III and III (Supplementary Figure 4b). VSD_I_, VSD_III_, and VSD_IV_ in the activated state and VSD_II_ in the resting state were also determined in both Ca_V_2.3 and Ca_V_2.2 (Supplementary Figure 4a–4b). However, a structural discrepancy was visualized at the ECL_IV_ between the two structures. The ECL_IV_ of Ca_V_2.3 extends from the pore domain and lies above the S1-S2_III_ linker, whereas the ECL_IV_ of Ca_V_2.2 is much shorter and wraps around the pore domain before touching the extracellular side of VSD_III_ (Figure 2a). Four residues on ECL_IV_ of Ca_V_2.3, namely, P1680, D1681, T1682, and T1683, are involved in the interactions with the residues on the S1-S2_III_ linker, especially V1176, L1177, T1178, and N1179, which consequently stitches the voltage sensing domain to the pore domain at the extracellular side (Figure 2a). To explore the functional roles of this interaction, we substituted ^1680^PDTT^1683^ on ECL_IV_ with four glycines (Ca_V_2.3^4G^) to disrupt the contacts between ECL_IV_ and S1-S2_III_ loop. Electrophysiological studies indicated that the voltage dependency of the activation curve of Ca_V_2.3^4G^ displayed a ∼5-mV positive shift (P < 0.0001, two-tailed unpaired *t*-test) compared to that of wild-type Ca_V_2.3 (Figure 2b and Supplementary Figure 5). We thus speculate that the interactions between ECL_IV_ and the S1-S2_III_ loop may stabilize the VSD_III_ in a certain conformation relative to the pore domain that requires less electrical energy to activate the channel, reminiscent of the cholesterol regulation on Ca_V_ channels, potentially by stabilizing interactions between the extracellular end of the S1-S2 helix hairpin and the pore domain^37,38^.

**Figure 2.**
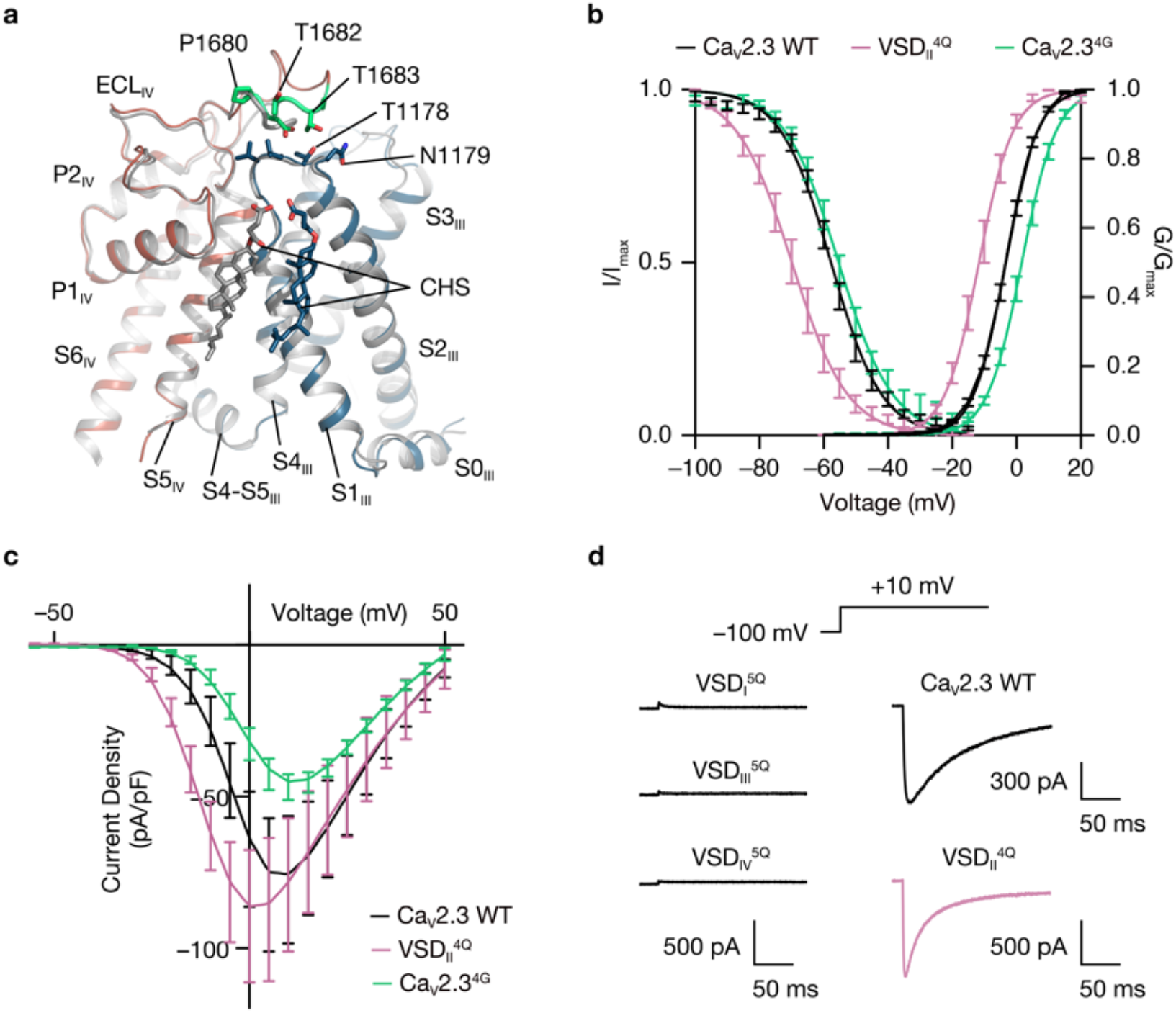
Voltage-sensing domains of the Ca_V_2.3 complex. **a**. Superimposition of the VSDs (DIII) and the pore domains (DIV) of the Ca_V_2.3 (red and blue) and Ca_V_2.2 (gray) demonstrating the interaction between the ECL_IV_ and the VSD_III_. Residues on ECL_IV_ in close contact with VSD_III_ (^1680^PDTT^1683^) are shown in sticks and colored in green. Cholesteryl hemisuccinate (CHS) molecules are shown as sticks and labeled. **b**. Steady-state activation and inactivation curves of the wild-type (WT) Ca_V_2.3 (black), VSD_II_^4Q^ (pink), and Ca_V_2.3^4G^ (green). To examine the voltage dependence of activation, HEK 293-T cells expressing the Ca_V_2.3 complex were tested by 200-ms depolarizing pulses between –60 and 50 mV from a holding potential of –100 mV, in 10-mV increments. To determination of inactivation curves, cells were stepped from a holding potential of –100 mV to pre-pulse potentials between –100 and –15 mV in 5-mV increments for 10 s. Activation curve, Ca_V_2.3 WT, n = 14; VSD_II_^4Q^, n = 7; Ca_V_2.3^4G^, n = 11. Steady-state inactivation curve, Ca_V_2.3 WT, n = 9; VSD_II_^4Q^, n = 7; Ca_V_2.3^4G^, n = 9. **c**. Current density of the wild-type Ca_V_2.3 and VSD_II_^4Q^. Ca_V_2.3 WT, n = 8; VSD_II_^4Q^, n = 8; Ca_V_2.3^4G^, n = 7. **d**. Voltage-clamp protocol and representative traces of the Ca_V_2.3 WT, VSD_I_^5Q^, VSD_II_^4Q^, VSD_III_^5Q^, and VSD_IV_^5Q^, elicited by a 200-ms test pulse at +10 mV.

The VSD_II_s of Ca_V_2.3 and Ca_V_2.2 were determined in the resting (S4 down) state, while most other VSDs in the voltage-gated channels, such as Ca_V_1.1, Ca_V_3.1, and Na_V_s, were determined in the activated (S4 up) state (Supplementary Figure 4c–4d). This finding suggests that VSD_II_ plays a unique role in the gating mechanism among Ca_V_2 members. Structural analyses of Ca_V_2.2 have suggested that the VSD_II_ is trapped in the resting state by a PIP_2_ molecule^27,28^. The cryo-EM map of Ca_V_2.3 is of high quality around S4-S5_II_ (Supplementary Figure 2e), and we identified a density lying above the S4-S5_II_ helix that serves as a wedge to hinder the upward movement of the S4 helix. However, the density in Ca_V_2.3 appears to be a single strip-shaped density and does not look like a PIP_2_ molecule. To shed light on the functional roles of VSD_II_ in the gating mechanism of Ca_V_2.3, we constructed a gating charge neutralization mutant on the VSD_II_ (VSD_II4Q_, R572Q/R575Q/R578Q/K581Q). Interestingly, the neutralization mutation of the VSD_II_ (VSD_II4Q_) exhibits ∼9-mV left shifts in the voltage dependency of both activation and steady-state inactivation compared to the wild-type (Figure 2b and Supplementary Figure 5a–5b). Consequently, the current density-voltage curve of the VSD_II_^4Q^ mutant was also left shifted (Figure 2c). Moreover, we tested whether the VSD_II_^4Q^ mutant has a distinct CSI profile. The cumulative inactivation of this mutant in response to action potential (AP) trains is markedly enhanced (Supplementary Figure 5c and 5e). These results suggested that the conformation of VSD_II_ influences the gating of Ca_V_2.3. However, its OSI kinetics remain unaltered (Supplementary Figure 5d). In contrast, the gating charge-neutralized mutation in the VSD_I_, VSD_III_, and VSD_IV_ resulted in failure to mediate inward current (Figure 2d), in line with previous results showing that the VSD_I_, VSD_III_, and VSD_IV_ are important for gating of the closely-related Ca_V_2.2^39-41^, while the VSD_II_ is not necessary for channel activation by sensing the depolarization of membrane potential; instead, the VSD_II_ is crucial to modulating channel properties, such as CSI and voltage dependency of channel activation and inactivation.

### Molecular mechanism of closed-state inactivation

Preferential closed-state inactivation is a featured kinetic characteristic of neuronal voltage-gated calcium channels^14,28^. During the state-transition pathway in the activation of Ca_V_ channels, CSI occurs preferentially in a specific pre-open closed state and a voltage-dependent manner. CSI can be visualized by the cumulative inactivation in response to action potential (AP) trains, as reported in previous studies, which demonstrated that the peak current triggered by each AP shrank sequentially, suggesting that a substantial amount of the channels turn inactivated after the repolarization of an AP. CSI plays an important role in the orchestrated modulation mechanism of Ca_V_ channels and is of vital importance for the precise regulation of physiological processes such as neurotransmitter release and synapse plasticity. CSI is detected in all neuronal Ca_V_2 channels at distinct levels^14^. The R-type Ca_V_2.3 displayed a more prominent CSI than the N-type Ca_V_2.2, and both showed far more prominent CSI than the P/Q-type Ca_V_2.1^14^. Previous structural investigations on the N-type Ca_V_2.2 channel have revealed that a conserved W-helix is the structural determinant of CSI, and W768 from the W-helix on the DII-DIII linker functions as a lid blocking the pore and stabilizes the intracellular gate in its closed states by hydrophobic interactions. However, the molecular mechanisms underlying the CSI of Ca_V_s are still not fully understood, as the conserved W-helix is unable to explain the disparity of CSI mechanisms among the Ca_V_2.1, Ca_V_2.2, and Ca_V_2.3 channels. The W-helix determined in Ca_V_2.2 is also conserved and well resolved in Ca_V_2.3 (Figure 3a–3b). The W-helix in Ca_V_2.3 (^772^RHHMSVWEQRTSQLRKH^788^) is a positively charged short helix and positioned underneath the intracellular gate, with W778 inserting into the gate and forming extensive interactions with residues from surrounding gating helices (Figure 3a–3b). These structural observations are consistent with the W-helix of Ca_V_2.2. First, we designed the Ca_V_2.3^W/Q^ (W778Q) to disrupt interactions between the W-helix and intracellular gate (Figure 3c–3g and Supplementary Figure 6). It turns out that the Ca_V_2.3^W/Q^ exhibited a ∼8-mV positive shift on the steady-state inactivation curve (Figure 3d) and an alleviated cumulative inactivation in response to AP trains (Figure 3e and 3g) without affecting the voltage dependence of channel activation (Figure 3c), consistent with the observations in Ca_V_2.2^28^, suggesting that the W778 is important for CSI process of Ca_V_2.3 channel. Moreover, we speculate that the positively charged residues on the W-helix could putatively respond to the membrane potential change and may be important for initiation of the CSI process. To evaluate our speculations, we constructed the Ca_V_2.3^RK/A^ mutant by substituting the R781, R786 and K787 with alanine (R781A/R786A/K787A). The activation curve of the Ca_V_2.3^RK/A^ mutant remains unaltered (Figure 3c). However, the CSI of Ca_V_2.3^RK/A^ mutant was significantly suppressed, exhibiting a ∼9 mV right shift on the steady-state inactivation curve (Figure 3d), an alleviated cumulative inactivation to AP trains (Figure 3e and 3g), and an accelerated recovery rate from CSI (Figure 3f and Supplementary Figure 6b). These alterations on the CSI profile suggested that the positively charged R781, R786 and K787 are critical for the CSI mechanism of Ca_V_2.3.

**Figure 3.**
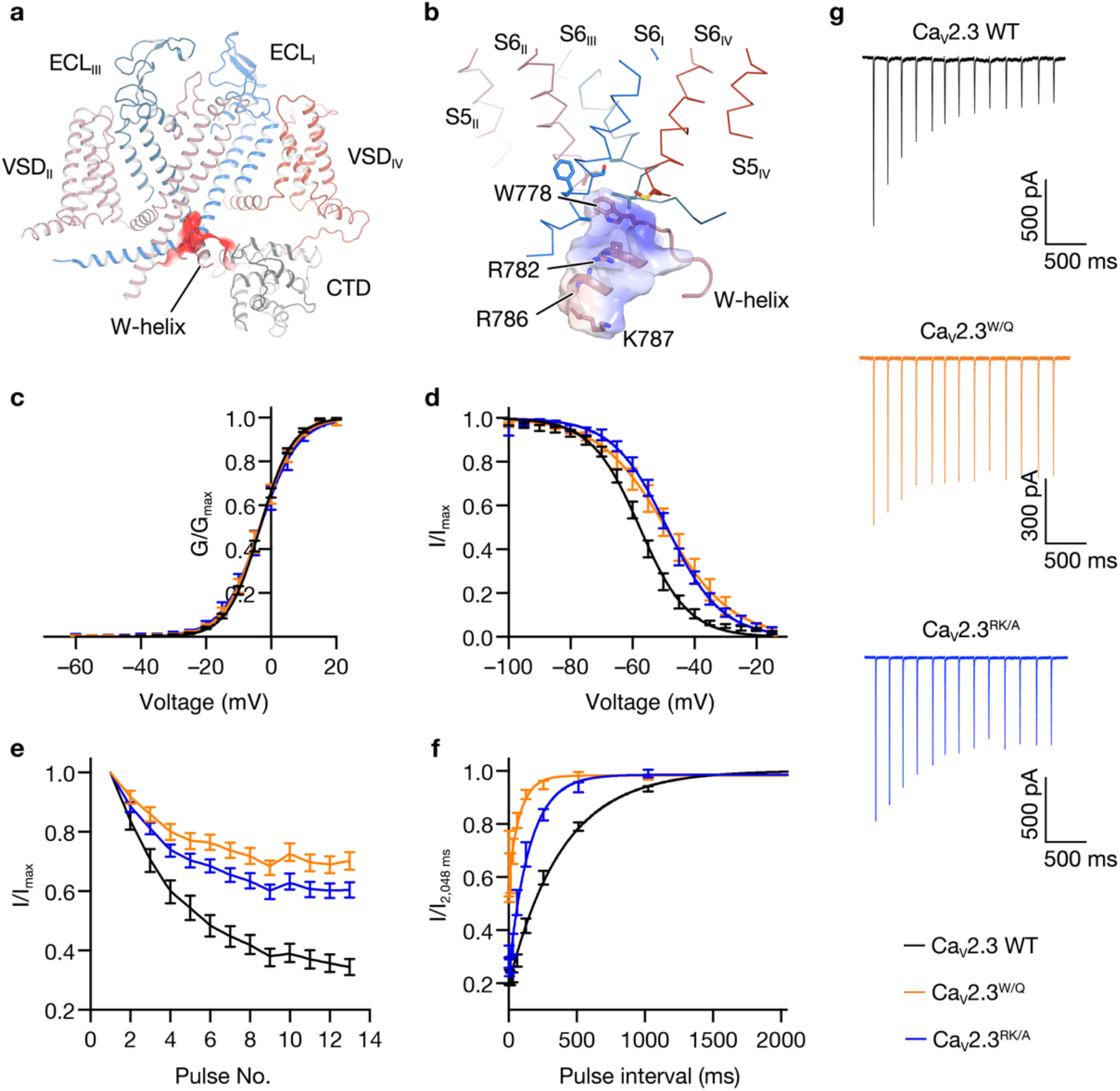
Closed-state inactivation mediated by the W-helix. **a**. Binding pocket of the W-helix. The W-helix is accommodated in a negatively charged pocket in the intracellular side of Ca_V_2.3. The Ca_V_2.3 α1E subunit is shown in cartoon, and the negatively charged pocket accommodating W-helix is shown as red surface. **b**. Zoomed-in view of the W-helix, which is stabilized in the intracellular gate at closed state. The W-helix is shown in cartoon and overlaid with an electrostatic surface. W778 and other residues involved in the hydrophobic or charge interactions are shown in sticks. **c**. Activation curves of the wild-type (WT) Ca_V_2.3 and the mutants. Ca_V_2.3 WT, n = 14; Δ*w-helix*, n = 7; Ca_V_2.3^RK/A^, n = 7. **d**. Steady-state inactivation curves of Ca_V_2.3 WT and the mutants. Ca_V_2.3 WT, n = 9; Δ*w-helix*, n = 8; Ca_V_2.3^RK/A^, n = 10. **e**. Inactivation ratio quantified using the current density (I) elicited by each spike of AP trains divided by the maximum current (I_Peak_) elicited by the first spike. Ca_V_2.3 WT, n = 13; Ca_V_2.3^W/Q^, n = 15; Δ*w-helix*, n = 9; Ca_V_2.3^RK/A^, n = 14. **f**. Recovery rate from the CSI, quantified by the inactivation ratio between the two peak currents (I and I_2,048ms_) obtained in the two-pulse protocol. HEK 293-T cells were held at –40 mV for 1,500 ms and were then stepped to –100 mV for a series of time intervals (4–2,048 ms) before a +10 mV test pulse (35 ms). Ca_V_2.3 WT, n = 6; Ca_V_2.3^W/Q^, n = 6; Δ*w-helix*, n = 6; Ca_V_2.3^RK/A^, n = 7. **g**. Representative current responses to the AP trains. The AP trains used to stimulate the HEK 293-T cells were recorded from a mouse hippocampal CA1 pyramidal neuron after current injection in whole-cell current-clamp mode. See Supplementary Figure 5 for voltage-clamp protocols and Method section for literature reference. Ca_V_2.3 WT, black; Ca_V_2.3^W/Q^, orange; Ca_V_2.3^RK/A^, blue.

Intriguingly, sequence alignment among the neuronal Ca_V_s revealed that a peptide segment (pre-W-helix) that resembled the W-helix is located adjacent to the W-helix of Ca_V_2.3 (Figure 4a). The sequence of pre-W-helix (^753^RHHMSMWEPRSSHLRER^769^) is nearly identical to that of the W-helix (∼65% identity), including the conserved tryptophan plug (W759) and positively charged residues (R762, R767, and R769) (Figure 4a). However, we noticed that a non-conserved proline (P761) was in the middle of the pre-W-helix, which may undermine the stability of both the pre-W-helix itself and its interactions with the gate. Moreover, in our cryo-EM map of the Ca_V_2.3 complex, the helical density beneath the gate perfectly fits the atomic model of the W-helix, enabling us to unambiguously determine that the W-helix, instead of the pre-W-helix, exists in our structure (Supplementary Figure 2e). Considering that the pre-W-helix is located immediately before the W-helix and that they share high sequence identity, we speculate that the pre-W-helix participates in the CSI event of Ca_V_2.3. To evaluate the contribution of the pre-W-helix to the CSI of Ca_V_2.3, we constructed two mutants by deleting the W-helix (Δ*w-helix*) and pre-W-helix (Δ*pre-w-helix*) (Figure 4b–4e, 4k, and Supplementary Figure 6). The Δ*w-helix* exhibited a ∼4-mV positive shift on the steady-state inactivation curve (Figure 4c) and alleviated cumulative inactivation in response to AP trains (Figure 4d and 4k) without affecting the voltage dependence of channel activation (Figure 4b), indicating that the W-helix plays pivotal roles in the CSI of Ca_V_2.3. However, compared with the CSI of Ca_V_2.2, which was almost abolished by deleting the W-helix, a substantial portion of the CSI in the Δ*w-helix* mutant remained unaltered (Figure 4d and 4k), suggesting that the CSI modulation of Ca_V_2.3 is distinct from that of Ca_V_2.2 and that other elements may also contribute to the CSI of Ca_V_2.3. We also used a two-pulse protocol to assess the recovery rate from CSI, i.e., the release process of CSI (Supplementary Figure 6a). Consistent with the electrophysiological results above, an accelerated recovery rate from CSI was observed in the Δ*w-helix* compared to the WT (Figure 4e and Supplementary Figure 6b). Strikingly, a negative-shift of ∼8 mV was detected on the inactivation curve of the Δ*pre-w-helix* mutant (Figure 4c), and its cumulative inactivation to AP trains was surprisingly enhanced (Figure 4d and 4k), demonstrating that the development process of CSI in the Δ*pre-w-helix* was significantly boosted. Considering that the pre-W and W helices are close to each other and share high sequence identity, we speculate that the pre-W-helix may serve as a competitive negative regulator to interfere with the binding of the intracellular gate to the W-helix (Figure 4f). In the absence of the pre-W-helix, the CSI is consequently enhanced (Figure 4c–4d, and 4k). Paradoxically, compared with the WT, the recovery rate from CSI of the Δ*pre-w-helix* mutant was substantially accelerated (Figure 4e and Supplementary Figure 6b). This suggests that once CSI occurs, the pre-W-helix appears to stabilize the W-helix for interaction with the gate during membrane potential repolarization, thereby slowing the recovery of Ca_V_2.3 from the CSI. We also designed a mutant by deleting both the pre-W-helix and the W-helix (Δ*pre-w*/Δ*w-helix*). This mutant displayed a ∼5-mV positive shift on the inactivation curve (Figure 4c), reduced cumulative inactivation to AP trains (Figure 4d and 4k), and an accelerated recovery rate from CSI (Figure 4e and Supplementary Figure 6b). These effects of this double deletion construct are essentially identical to those of the Δ*w-helix*, demonstrating that the modulatory role of the pre-W-helix on CSI is largely dependent on the W-helix, probably by regulating the binding or dissociation of the W-helix with the intracellular gate. To further investigate the regulatory mechanism on the CSI of Ca_V_2.3, we constructed two mutants by neutralizing the arginines (Ca_V_2.3^preR/Q^, R753Q/R762Q/R767Q/R769Q) or substituting the tryptophan on the pre-W-helix with glutamine (Ca_V_2.3^preW/Q^, W759Q) (Figure 4g–4k and Supplementary Figure 6). Impressively, the mutants Ca_V_2.3^preR/Q^ and Ca_V_2.3^preW/Q^ exhibit similar gating kinetics and voltage dependence of channel activation and inactivation, but significantly different from that of the WT Ca_V_2.3 channel (Figure 4g–4k, Supplementary Figure 5a–5b and Supplementary Figure 6b). In particular, they displayed a ∼7-mV negative shift on inactivation curve (Figure 4h), enhanced cumulative inactivation to AP trains (Figure 4i and 4k), and an accelerated recovery rate from CSI (Figure 4j and Supplementary Figure 6b), highly identical to the CSI profile of the Δ*pre-w-helix*. The kinetic characteristics of these mutants indicated that the positively charged residues and W759 are important for the regulatory effect of the pre-W-helix. Our data also show that the development and release of CSI are two independent processes. In particular, the Δ*pre-w-helix*, Ca_V_2.3^preR/Q^ and Ca_V_2.3^preW/Q^ exhibited enhanced cumulative inactivation during AP trains (Figure 4d, 4i and 4k) but a faster recovery rate from CSI (Figure 4e, 4j and Supplementary Figure 6b) at a membrane potential of –100 mV, suggesting that the channel may have distinct conformational states along the activation pathway. The W-helix may bind preferentially to the channel in the intermediate closed state(s) near the open state, thus leading to the largest inactivation before the channel opens (Figure 4f). At strongly hyperpolarized potentials (∼100 mV), the channel is probably stabilized in a different closed state, exhibiting a lower affinity to the W-helix, causing the W-helix to detach from the gate and the channel to recover from the CSI (Figure 4f). Taken together, the pre-W-helix in Ca_V_2.3 plays significant roles in regulating the association or dissociation of the W-helix with the intracellular gate and thus exerts a regulatory effect on the development and release of the CSI.

**Figure 4.**
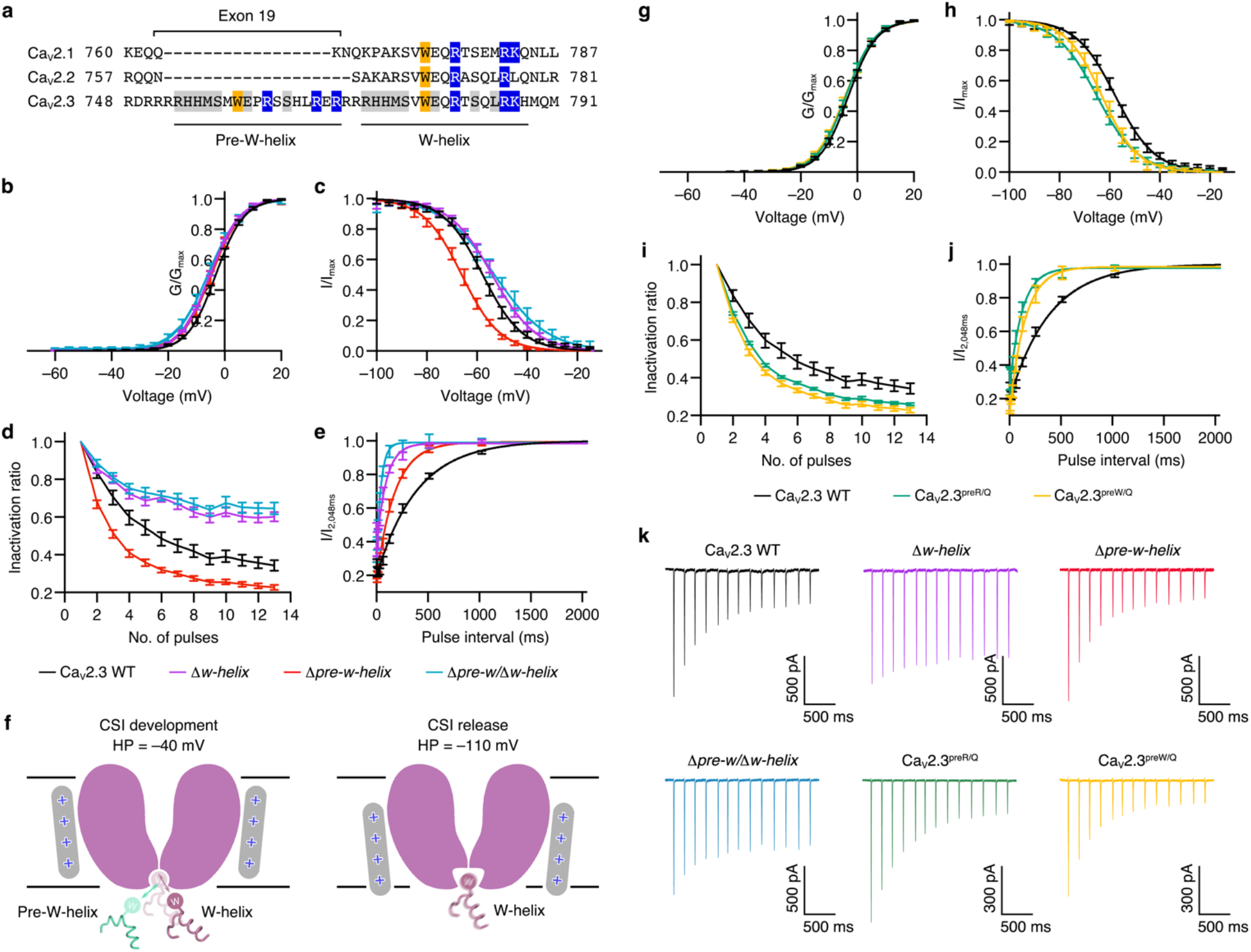
CSI regulation by pre-W-helix. **a**. Sequence alignment of the DII-III linker among the Ca_V_2 members. Residues on the pre-W-helix and W-helix are underlined and labeled, respectively. Exon 19 on the genomic DNA of Ca_V_2.3 encoding the pre-W-helix is also labeled. **b** and **g**. Activation curves of the wild-type Ca_V_2.3 and the mutants. Ca_V_2.3 WT, n = 14; Δ*w-helix*, n = 7; Δ*pre-w-helix*, n = 7; Δ*pre-w/Δw-helix*, n = 7; Ca_V_2.3^preR/Q^, n = 7; Ca_V_2.3^preW/Q^, n = 8. **c** and **h**. Steady-state inactivation curves of wild-type Ca_V_2.3 and the mutants. Ca_V_2.3 WT, n = 9; Δ*w-helix*, n = 8; Δ*pre-w-helix*, n = 7; Δ*pre-w/Δw-helix*, n = 7; Ca_V_2.3^preR/Q^, n = 8; Ca_V_2.3^preW/Q^, n = 6. **d** and **i**. Inactivation ratio quantified using I/I_Peak_ elicited by the AP trains. Ca_V_2.3 WT, n = 13; Δ*w-helix*, n = 9; Δ*pre-w-helix*, n = 13; Δ*pre-w/Δw-helix*, n = 14; Ca_V_2.3^preR/Q^, n = 9; Ca_V_2.3^preW/Q^, n = 13. **e** and **j**. Recovery rate from the CSI, quantified using the previously described two-pulse protocol. Ca_V_2.3 WT, n = 6; Δ*w-helix*, n = 6; Δ*pre-w-helix*, n = 7; Δ*pre-w/Δw-helix*, n = 6; Ca_V_2.3^preR/Q^, n = 8; Ca_V_2.3^preW/Q^, n = 7. **f**. Proposed model of the putative regulatory role of the pre-W-helix in the CSI of Ca_V_2.3. Membrane planes are indicated using grey lines. Pore domains are depicted as cartoon and colored in purple. Positively-charged S4 helices are shown as grey rounded bars. Pre-W-helix and the W-helix are shown as cartoon. In the development process of CSI (left), the intracellular gate might have a high affinity to W-helix. As a consequence, W-helix and pre-W-helix competitively bind to the gate. In the release process of CSI (right), the gate might adopt an alternative conformation and the W-helix is likely to disassociate from the gate. **k**. Representative current responses stimulated by AP trains. See Supplementary Figure 5 for voltage-clamp protocols and Method section for literature reference. Ca_V_2.3 WT, black; Δ*w-helix*, purple; Δ*pre-w-helix*, red; Δ*pre-w/Δw-helix*, green-cyan; Ca_V_2.3^preR/Q^, green; Ca_V_2.3^preW/Q^, yellow.

Interestingly, on the genomic DNA of Ca_V_2.3, the pre-W-helix is encoded by exon 19 (residues R748–R769) (Figure 4a). Early studies have reported that alternative splicing events might occur at this site, as exon 19 is spliced out in the mature mRNA encoding Ca_V_2.3e, an isoform of Ca_V_2.3 that displays decreased Ca^2+^ sensitivity in its calcium-dependent modulation mechanisms^42^. Gene expression profiling has revealed that Ca_V_2.3e is enriched in endocrine tissues, including the kidney and pancreas, and the majority of Ca_V_2.3 in the brain contains the pre-W-helix^43^. This implies that the kinetic variability of Ca_V_2.3 mediated by the pre-W-helix is an important regulatory mechanism to fine-tune the properties of the Ca_V_2.3 channel to adapt to the distinct physiological needs of neuronal and endocrinal excitable cells^42^.

### Modulation of open-state inactivation

Upon the opening of the pore, Ca_V_ channels undergo an inactivation process called open-state inactivation (OSI), which describes the mechanism by which the intracellular gate shifts swiftly from the open state into the inactivation state^33,44^. This process is an intrinsic mechanism that precisely regulates calcium influx into cells during depolarization. Neuronal (P/Q-, N- and R-type) Ca_V_2 channels bear a stronger OSI and mediate a rapidly inactivated current, while cardiac (L-type) Ca_V_ channels display a much weaker OSI, mediating a long-lasting current^45^. Dysfunction of OSI, i.e., the gain-of-function of neuronal Ca_V_s, is linked to a series of neurological disorders, including trigeminal neuralgia^46^, myoclonus-dystonia-like syndrome^47^, and epileptic encephalopathies^21,24^. Although the structures of L-type Ca_V_1.1^25^ and N-type Ca_V_2.2^27,28^ complexes have been elucidated at high resolution, the structural basis for their distinct OSI properties remains elusive. Intriguingly, when comparing the EM density maps of the Ca_V_1.1, Ca_V_2.2, and Ca_V_2.3 complexes, we observed that AID displayed a blurred density in Ca_V_1.1 but was clearly resolved in Ca_V_2.2 and Ca_V_2.3 (Supplementary Figure 7). Previous studies suggested that the AID helix is able to regulate the OSI in Ca_V_ channels^33,34^.

In the structure of Ca_V_2.3, the S6_II_ helix is much longer than that of Ca_V_1.1 and extends into the cytosol. Interestingly, we identified a negatively charged domain ^715^DEQEEEE^721^ in the intracellular juxtamembrane region of the S6_II_ helix (S6_IINCD_) (Figure 5a–5b). Taking a closer look at the structure, this negatively charged region forms electrostatic interactions with the positively charged R590 on S4-S5_II_, as well as R371 and R378 on the AID (Figure 5b). Strikingly, the S6_IINCD_ is conserved among P/Q-, N- and R-type channels (Figure 5c). The equivalent segment of L-type Ca_V_ channels contains fewer negatively charged residues, interspersed with some positively charged residues, indicating that the electrostatic interactions between S6_IINCD_ and AID are not present in the L-type Ca_V_ channels, which is consistent with structural observations that the AID of Ca_V_1.1 has high motility (Figure 5c and Supplementary Figure 7). Additionally, the positively charged R590 and R378 are conserved only in Ca_V_2 channels. R378 is further reverted to a negatively charged glutamate in the Ca_V_1 subfamily, reflecting the coevolutionary linkages within this interaction site (Figure 5c). We speculate that the charge interactions centered on S6_II_^NCD^ is critical to regulating the OSI of Ca_V_2 channels.

**Figure 5.**
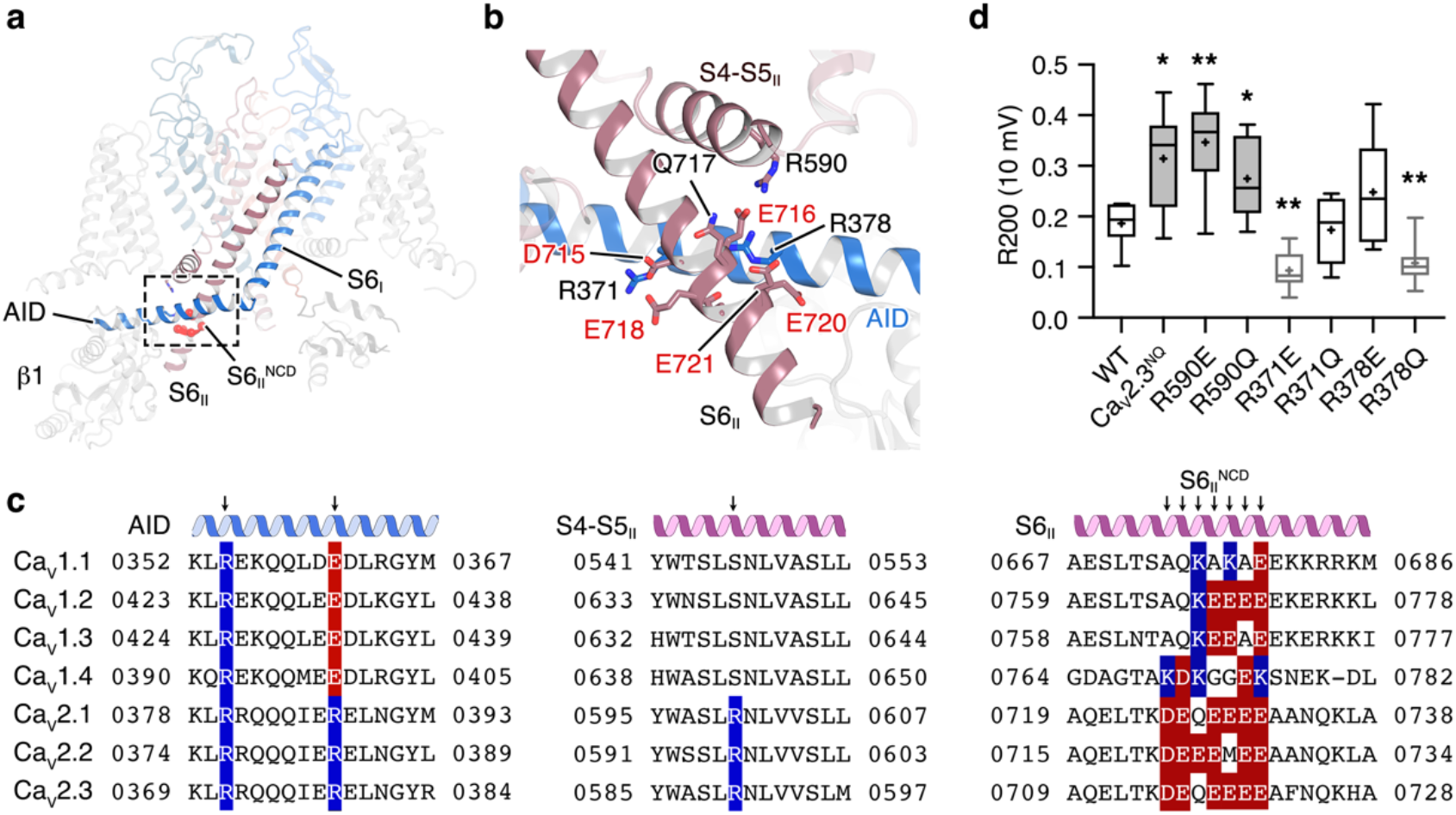
Structural basis of the modulation of open-state inactivation. **a**. Interaction of the AID between the negatively-charged domain on S6_II_ (S6_II_^NCD^) and S4-S5_II_. The Ca_V_2.3 α1E subunit is shown as cartoon. The S6_II_^NCD^ is overlaid as red spheres. **b**. Zoomed-in view of the side-chain interactions among S6_II_^NCD^, AID, and S4-S5_II_. The S6_II_^NCD^ (^715^DEQEEEE^721^) is shown as sticks. Residues involved in charge interactions on AID and S4-S5_II_ are also shown as sticks. Negatively charged residues within the S6_II_^NCD^ are highlighted using red labels. **c**. Sequence alignment of the interaction sites among Ca_V_1 and Ca_V_2 members. Secondary structures are labeled over the sequences. Mutation sites are indicated by arrows. Residues with positively- and negatively-charged side chains are highlighted by blue and red, respectively. **d**. Ratio of open-state inactivation (R200) at 10-mV test pulses, measured using the mean current at the end of the 200-ms test pulse divided by the peak amplitude. WT, n = 6; Ca_V_2.3^NQ^, n = 10; R590E, n = 6; R590Q, n = 10; R378E, n = 6; R378Q, n = 9; R371E, n = 6; R371Q, n = 9. Data are plotted as a box plot. The box encompasses the interquartile range (25th–75th percentile). Mean values and medians are indicated using plus signs and dashes, respectively. Significances were determined using two-sided, unpaired *t*-test. P values, Ca_V_2.3 WT vs. mutants; 0.01 (Ca_V_2.3^NQ^), 0.005 (R590E), 0.02 (R590Q), 0.004 (R371E), and 0.004 (R378Q).

To validate our hypothesis, we mutated the key residues to disrupt electrostatic interactions cross-linking the S6_II_, S4-S5 and AID helices, including replacing ^715^DEQEEEE^721^ with ^715^NNQNNNN^721^ (Ca_V_2.3^NQ^), R590Q, R378Q/E, and R371Q/E (Figure 5d and Supplementary Figure 8). We employed Ba^2+^ as the charge carrier in our whole-cell patch clamp analysis to exclude the effects of calcium-dependent inactivation (CDI) of the Ca_V_2.3 channels. The Ca_V_2.3^NQ^ mutant mediates a current that decays much more slowly than the wild-type Ca_V_2.3 during a 200-ms test pulse (Supplementary Figure 8a–8b), suggesting that OSI is remarkably decreased. To quantify the OSI of the Ca_V_2.3 mutants, we employed the R200 value as an indicator, which is calculated by the mean current density at the end of the 200-ms test pulse divided by the peak amplitude (Figure 5d and Supplementary Figure 8c). Specifically, the mean value of R200 increased from 0.17±0.01 in the wild-type Ca_V_2.3 to 0.31±0.03 in Ca_V_2.3^NQ^ under the test pulse holding at 10 mV (Figure 5d and Supplementary Figure 8c). Moreover, R590E and R590Q also exhibited remarkably suppressed OSI, with increased R200 values of 0.35±0.04 and 0.27±0.03, respectively (10-mV test pulse) (Figure 5d and Supplementary Figure 8c), suggesting that the R590-S6_IINCD_ interaction plays an essential role in the development of OSI in Ca_V_2 channels. In contrast, mutants on the AID side show complicated effects on OSI kinetics (Figure 5d and Supplementary Figure 8c). In particular, R378Q displayed an enhanced OSI, with a decreased R200 value of 0.11±0.02 (10-mV test pulse). R371E, which is located in the adjacent region of R378, exhibited an enhanced OSI as well, displaying an R200 value of 0.09±0.01 (10-mV test pulses). Nevertheless, OSI of the R371Q and R378E mutants do not show significant difference with that of the WT Ca_V_2.3 channel (Figure 5d and Supplementary Figure 8c). Moreover, the time course of channel inactivation could be well fitted by a single exponential. Conclusions drawn using time constants are nearly identical to those using R200 values (Supplementary Figure 8d), further supporting that S6_IINCD_, R371 (AID), R378 (AID), and R590 (S4-S5_II_) play important roles in OSI modulation. However, the interaction between the AID and S6_IINCD_ could go beyond the current structures of Ca_V_ channels and is worth investigation in future studies.

## Methods

### Expression and protein purification of the human Ca_V_2.3 complex

Full-length Ca_V_2.3 α1E (CACNA1E, UniProt ID: Q15878 isoform 1, or Ca_V_2.3d), α2δ1 (CACNA2D1, UniProt ID: P54289), and β1 (CACB1, UniProt ID: Q02641) were amplified from a human cDNA library and subcloned into pEG BacMam vectors. To detect the expression and assembly levels of the Ca_V_2.3 complex, a superfolder GFP, an mCherry, and an mKalama tag were fused to the C-terminal, N-terminal, and N-terminal regions of the Ca_V_2.3 α1, α2δ1, and β1 subunits, respectively. Twin-Strep tags were tandemly inserted into the C-terminal and N-terminal of the α1 and α2δ1 subunits, respectively. The Bac-to-Bac baculovirus system (Invitrogen, USA) was used to conduct protein expression in HEK 293-F cells (Gibco, USA) following the manufacturer’s protocol. The bacmids were prepared using DH10Bac competent cells, and P1 viruses were generated from *Sf*9 cells (Invitrogen, USA) after bacmid transfection. P2 viruses (1%, v/v) were used to infect HEK 293-F cells supplemented with 1% (v/v) fetal bovine serum. The cells were cultured at 37°C and 5% CO_2_ for 12 hours before the addition of 10 mM sodium butyrate to the medium. The cells were cultured at 30°C and 5% CO_2_ for another 48 h before harvest. No authentication was performed for the HEK 293-F or the *Sf* 9 cell line. No *Mycoplasma* contamination was observed.

The cell pellets were resuspended at 4°C using Buffer W containing 20 mM HEPES pH 7.5, 150 mM NaCl, 5 mM β-mercaptoethanol (β-ME), 2 μg/mL aprotinin, 1.4 μg/mL leupeptin, and 0.5 μg/mL pepstatin A by a Dounce homogenizer, followed by centrifugation at 36,900 rpm for 1 h to collect the membrane. The membrane was resuspended again using Buffer W and solubilized by the addition of 1% (w/v) n-dodecyl-β-D-maltoside (DDM) (Anatrace, USA), 0.15% (w/v) cholesteryl hemisuccinate (CHS) (Anatrace, USA), 2 mM adenosine triphosphate (ATP) and 5 mM MgCl_2_ on a rotating mixer at 4°C for 2 h. Addition of ATP and MgCl_2_ is to remove associated heat shock proteins. The insoluble debris of cells was removed by another centrifugation at 36,900 rpm for 1 h. The supernatant was passed through a 0.22 μm filter (Millipore, USA) before being loaded into a 6 mL Strep-Tactin Superflow high capacity resin (IBA Lifesciences, USA). The resin was washed using 6 column volumes of Buffer W1 (20 mM HEPES pH 7.5, 150 mM NaCl, 5 mM β-ME, 0.03% (w/v) glycol-diosgenin (GDN) (Anatrace, USA), 2 mM ATP, and 5 mM MgCl_2_). The purified Ca_V_2.3 complex was eluted using 15 mL elution buffer (20 mM HEPES pH 7.5, 150 mM NaCl, 5 mM β-ME, 0.03% (w/v) GDN, and 5 mM d-Desthiobiotin (Sigma-Aldrich, USA)) and concentrated to 1 mL using a 100 kDa MWCO Amicon (Millipore, USA). The concentrated protein sample was subjected to further size-exclusion chromatography (SEC) by a Superose 6 Increase 10/300 GL gel filtration column (GE Healthcare, USA) using a flow rate of 0.3 mL/min and a running buffer containing 20 mM HEPES pH 7.5, 1.5 mM NaCl, 5 mM β-ME, and 0.01% (w/v) GDN (Anatrace, USA). The monodispersed peak fraction within 12–13.5 mL was pooled and concentrated to 5 mg/mL before preparing the cryo-EM grids.

### Cryo-EM sample preparation and data collection

Holey carbon grids (Au R1.2/1.3 300 mesh) (Quantifoil Micro Tools, Germany) were glow-discharged using H_2_ and O_2_ for 60 s before being loaded with 2.5 μL purified Ca_V_2.3 complex. The grids were automatically blotted for 4 s at 4°C and 100% humidity and flash-frozen in liquid ethane using a Vitrobot Mark IV (Thermo Fisher Scientific, USA). Cryo-EM data were collected using a 300 kV Titan Krios G2 (Thermo Fisher Scientific, USA) equipped with a K2 Summit direct electron detector (Gatan, USA) and a GIF Quantum LS energy filter (Gatan, USA). The dose rate was set to ∼9.2 e^−^/(pixel*s), and the energy filter slit width was set to 20 eV. A total exposure time of 6.72 s was dose-fractioned into 32 frames. A nominal magnification of ×130,000 was used, resulting in a calibrated super-resolution pixel size of 0.52 Å on images. SerialEM^48^ was used to automatically acquire the movie stacks. The nominal defocus range was set from –1.2 μm to –2.2 μm.

### Cryo-EM data processing

Motion correction was performed on 2,096 movie stacks using MotionCor2^49^ with 5 × 5 patches, generating dose-weighted micrographs. The parameters of the contrast transfer function (CTF) were estimated using Gctf^50^. Particles were initially picked using the blob picker in cryoSPARC^51^, followed by 2D classifications to produce 2D templates and *ab initio* reconstruction to generate an initial reference map. Another round of particle picking was conducted using Template Picker in cryoSPARC, generating a dataset of 787,518 particles that were used for further processing. All data processing steps were performed in RELION-3.1_52_ unless otherwise specified. A round of multi-reference 3D classification was conducted against one good and 4 biased references, generating 5 classes. Class 5 (75.2%), which was calculated using the good reference, displayed a classical shape of Ca_V_ complexes featuring a transmembrane subunit and two soluble subunits residing on both sides of the micelle. Particles from class 5 were re-extracted and subjected to another round of 3D classification, resulting in 6 classes. Classes 1, 4, and 5 (43.5%) displayed well-resolved structural features, including continuous transmembrane helices of the α1E subunit and secondary structures within the α2δ1 and β1 subunits. To improve the map quality, Bayesian polish and CTF refinement were then conducted. The following 3D auto refinement generated a 3.1-Å map. The particle dataset was then imported back to cryoSPARC, where the final map was generated by Non-uniform (NU) refinement, which was reported at 3.1 Å according to the golden-standard *Fourier* shell correlation (GSFSC) criterion.

### Model building

The cryo-EM map of Ca_V_2.3 was reported at near-atomic resolution, which enabled us to reliably build and refine the model. The structure of the Ca_V_2.2-α2δ1-β1 complex (PDB ID: 7VFS)^28^ was selected as the starting model because of the high sequence identity and was docked into the map of Ca_V_2.3 complexes using UCSF Chimera^53^. Side chains of the a1 subunit were manually mutated according to the sequence alignment between Ca_V_2.3 and Ca_V_2.2 and adjusted according to the EM density using *Coot*^54^. Side chains of α2δ1 were also manually adjusted according to the EM density. β1 was initially fit into the EM maps as a rigid body and manually refined against a low-resolution map of the Ca_V_2.3 complex in *Coot* due to local structural heterogeneity. The manually adjusted models were then automatically refined against the cryo-EM maps using the integrated Real Space Refinement program within the PHENIX software package^55^. Model stereochemistry was also evaluated using the Comprehensive validation (cryo-EM) tool in PHENIX.

All the figures were prepared using Open-Source PyMOL (Schrödinger, USA), UCSF Chimera^53^, or UCSF ChimeraX^56^.

### Whole-cell voltage-clamp recordings of Ca_V_2.3 channels in HEK 293-T cells

HEK 293-T cells were cultured with Dulbecco’s modified Eagle’s medium (DMEM) (Gibco, USA) supplemented with 15% (v/v) fetal bovine serum (FBS) (PAN-Biotech, Germany) at 37°C with 5% CO_2_. The cells were grown in culture dishes (d = 3.5 cm) (Thermo Fisher Scientific, USA) for 24 h and then transiently transfected with 2 μg of control or mutant plasmid expressing the human R-type Ca_V_2.3 calcium channel complex (Ca_V_2.3 α1E, β1, α2δ1) using 1.2 μg of Lipofectamine 2000 Reagent (Thermo Fisher Scientific, USA). Patch-clamp experiments were performed 12 to 24 hours post-transfection at room temperature (21∼25°C) as described previously. Briefly, cells were placed on a glass chamber containing 105 mM NaCl, 10 mM BaCl_2_, 10 mM HEPES, 10 mM D-glucose, 30 mM TEA-Cl, 1 mM MgCl_2_, and 5 mM CsCl (pH = 7.3 with NaOH and an osmolarity of ∼310 mosmol^-1^). Whole-cell voltage-clamp recordings were made from isolated, GFP-positive cells using 1.5∼2.5 MΩfire-polished pipettes (Sutter Instrument, USA) filled with standard internal solution containing 135 mM K-gluconate, 10 mM HEPES, 5 mM EGTA, 2 mM MgCl_2_, 5 mM NaCl, and 4 mM Mg-ATP (pH = 7.2 with CsOH and an osmolarity of ∼295 mosmol^-1^). Whole-cell currents were recorded using an EPC-10 amplifier (HEKA Electronik, Germany) at a 20 kHz sample rate and were low-pass filtered at 5 kHz. The series resistance was 2∼4.5 MΩ and was compensated 80∼90%. The data were acquired by the PatchMaster program (HEKA Electronik, Germany).

To obtain activation curves of Ca_V_2.3 channels, cells were held at –100 mV, and then a series of 200-ms voltage steps from –60 mV to +50 mV in 5-mV increments were applied. The steady-state inactivation properties of Ca_V_2.3 channels were assessed with 10-s holding voltages ranging from –100 mV to –15 mV (5-mV increments) followed by a 135-ms test pulse at +10 mV. To assess the time-dependent recovery from CSI, cells were depolarized to –40 mV (pre-pulse) for 1500 ms to allow Ca_V_2.3 channels to enter CSI, and recovery hyperpolarization steps to –100 mV were applied for the indicated period (4 ms – 2,048 ms), followed by a 35-ms test pulse at +10 mV. To assess the cumulative inactivation of Ca_V_2.3 channels in response to AP trains, the cells were held at –100 mV, and then the physiologically relevant AP trains was applied. The AP trains used to stimulate the HEK 293-T cells were recorded from a mouse hippocampal CA1 pyramidal neuron after current injection in the whole-cell current-clamp mode^57^. The spike pattern contained 13 action potentials in 2 s (mean frequency = 6.5 Hz). The percentage inactivation of Ca_V_2.3 channels was calculated from the first spike eliciting maximal current to the other spikes in the AP trains. To analyze the extent of OSI, the ratio of remaining currents at 200 ms post-depolarization and the peak currents was calculated.

### Electrophysiological data analysis

All data are reported as the mean ± SEM. Data analyses were performed using Origin 2019b (OriginLab, USA) and Prism 9 (GraphPad, USA).

Steady-state activation curves were generated using a Boltzmann equation.

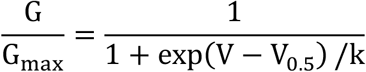

where G is the conductance, calculated by G = I/(V-V_rev_), where I is the current at the test potential and V_rev_ is the reversal potential; G_max_ is the maximal conductance of the Ca_V_2.3 channel during the test pulse; V is the test potential; V_0.5_ is the half-maximal activation potential; and k is the slope factor.

Steady-state activation and inactivation curves were generated using a Boltzmann equation.

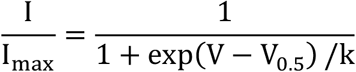

where I is the current at the indicated test pulse; I_max_ is the maximal current of Ca_V_2.3 activation during the test pulse; V is the test potential; V_0.5_ is the half-maximal inactivation potential; and k is the slope factor.

Recovery curves from CSI were calculated from the results of 7 to 9 independent experiments where a series of recovery traces from inactivation time points were acquired.

The data were fit using a single exponential of the following equation.

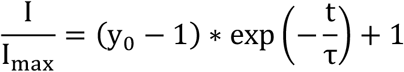

where I is the current at the indicated intervals; I_max_ is the current at 2048 ms; y_0_ is the remaining current at –40 mV for 1500 ms; t is the indicated hyperpolarization time; and τ is the time constant of recovery from CSI.

Statistical significance (*p* < 0.05) was determined using unpaired Student’s t tests or one-way ANOVA with Tukey’s *post hoc* test.

## Author contributions

Y.Z. conceived the project and supervised the research. Y.G. and Y.Q. carried out the molecular cloning experiments. Y.G. expressed and purified protein samples. Y.D. prepared samples for cryo-EM study. Y.G. and B.Z. carried out cryo-EM data collection. Y.G. processed the cryo-EM data. Y.G. built and refined the atomic model. Y.Z., Y.G., X.C.Z., and Y.W. analyzed the structure. Y.Z., Z.H., Y.G., S.X., and J.S. designed the electrophysiological experiments. S.X., X.C., H.X., C.P., and S.L. conducted the whole-cell voltage patch-clamp analysis. Y.G. wrote the original draft of the manuscript and prepared the figures. Y.Z., Y.G. and X.C.Z. edited the manuscript with input from all authors.

## Acknowledgments

We thank Xiaojun Huang, Xujing Li, Lihong Chen, and other staff members at the Center for Biological Imaging (CBI), Core Facilities for Protein Science at the Institute of Biophysics, Chinese Academy of Science for their support in cryo-EM data collection. We thank Yan Wu for his research assistance. This work is funded by the National Key Research and Development Program of China (Grant No. 2021YFA1301501 to Y.Z.), the Chinese Academy of Sciences Strategic Priority Research Program (Grant No. XDB37030304 to Y.Z.; Grant No. XDB37030301 to X.C.Z.), the National Natural Science Foundation of China (Grant No. 92157102 to Y.Z.; Grant No. 31971134 to X.C.Z.; Grant No. 81371432 to Z.H.), Chinese National Programs for Brain Science and Brain-like Intelligence Technology (Grant No. 2022ZD0205800 to Y.Z.; Grant No. 2021ZD0202102 to Z.H.).

## Data availability

The cryo-EM density map of the Ca_V_2.3-α2δ1-β1 complex has been deposited in the Electron Microscopy Data Bank (EMDB) under the accession code EMD-33285. The coordinate for the Ca_V_2.3 complex has been deposited in the Protein Data Bank (PDB) under the PDB ID 7XLQ.

## Conflict of interest

All authors declare that there is no conflict of interest that could be perceived as prejudicing the impartiality of the research reported.

## Notes

### Competing Interest Statement

The authors have declared no competing interest.

