## Supplementary Information for "Molecular insights into the gating mechanisms of voltage-gated calcium channel Ca_V_2.3"

This document contains Supplementary Figure 1–8 and Supplementary Table 1.

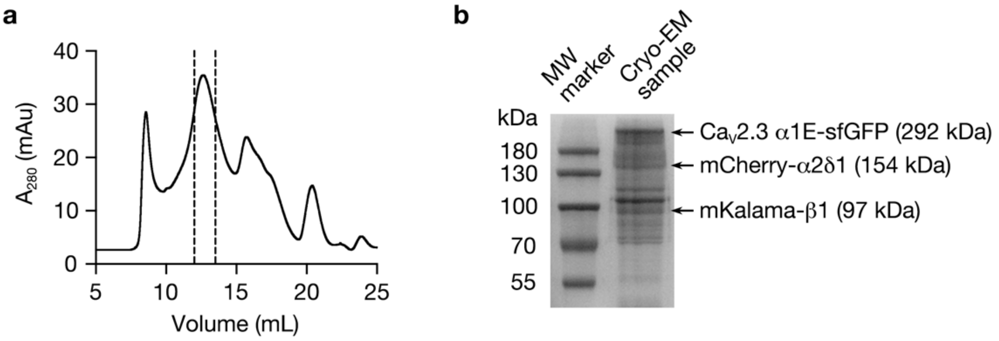


Supplementary Figure 1. Protein purification of the $Ca_V2.3$-α2δ1-β1 complex

**a**. Size-exclusion chromatogram of the purified protein sample of $Ca_V2.3$ complex. Peak fractions marked within the dashed lines were pooled and concentrated for the cryo-EM studies. **b**. Coomassie blue-stained SDS-PAGE gel of the purified $Ca_V2.3$ complex. Bands representing the $Ca_V2.3$ α1E, α2δ1, and β1 subunits were labeled. The experiments were repeated independently for more than 3 times with identical results.

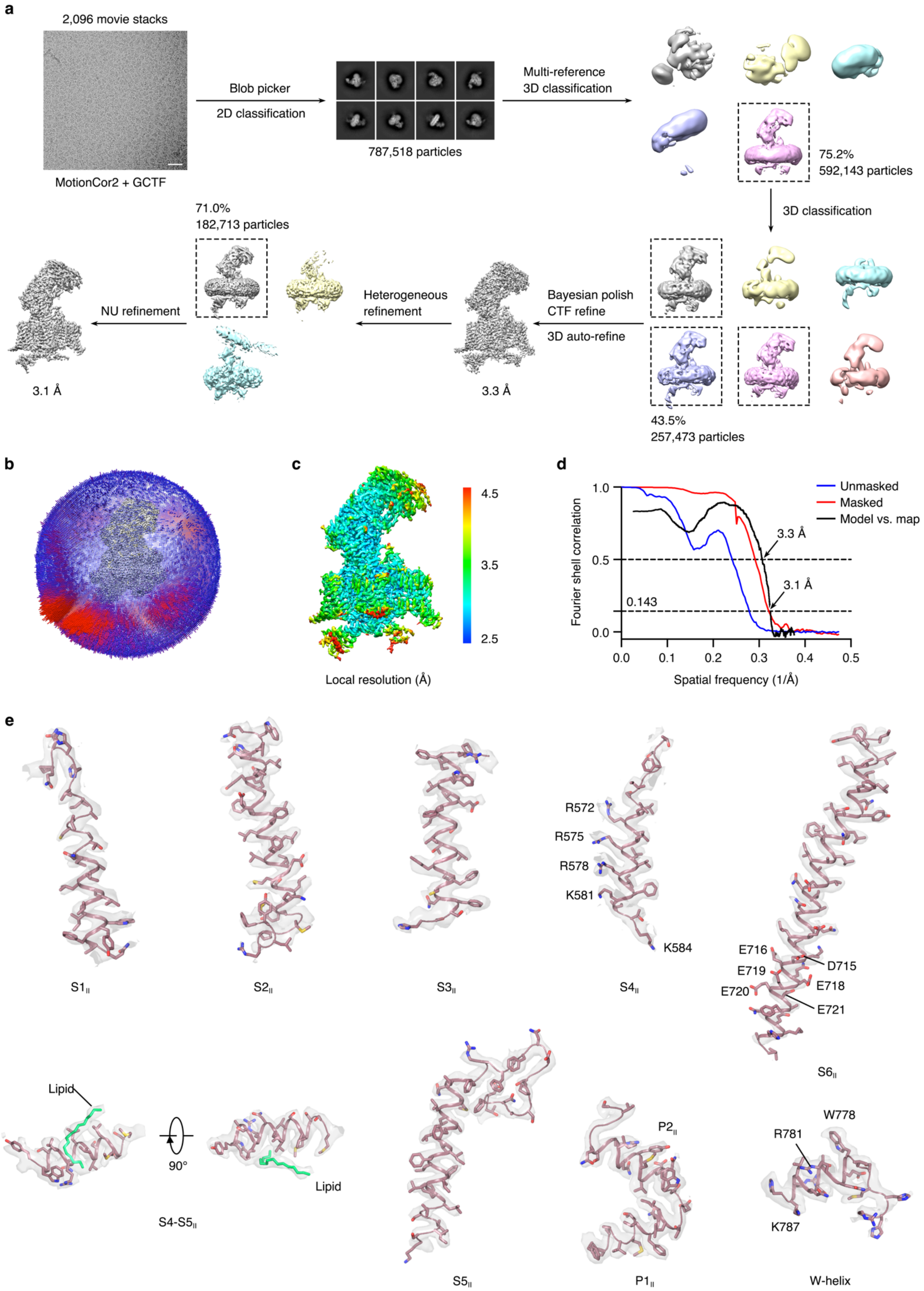

a
2,096 movie stacks
MotionCor2 + GCTF
Blob picker
2D classification
787,518 particles
Multi-reference
3D classification
75.2%
592,143 particles
3D classification
43.5%
257,473 particles
Bayesian polish
CTF refine
3D auto-refine
3.3 Å
Heterogeneous
refinement
71.0%
182,713 particles
NU refinement
3.1 Å
b
c
4.5
3.5
2.5
Local resolution (Å)
d
Fourier shell correlation
Spatial frequency (1/Å)
Unmasked
Masked
Model vs. map
3.3 Å
3.1 Å
0.143
1.0
0.5
0.0
0.0
0.1
0.2
0.3
0.4
0.5
e
S1II
S2II
S3II
R572
R575
R578
K581
K584
S4II
E716
D715
E719
E718
E720
E721
S6II
Lipid
90°
Lipid
S4-S5II
S5II
P2II
P1II
W778
R781
K787
W-helix

### Supplementary Figure 2. Cryo-EM data processing of the $Ca_V2.3$-α2δ1-β1 complex

**a**. Flow chart of cryo-EM data processing. A total of 787,518 particles were picked form 2,096 micrographs. A representative motion-corrected micrograph of the dataset was shown here (bar = 400 Å). A round of multi-reference 3D classification, and a further round of 3D classification against a single starting reference were performed to clean particles, followed by Bayesian polish and contrast transfer function (CTF) refinement to improve the map quality. Heterologous refinement and non-uniform (NU) refinement generated the final map, which was reported at 3.1 Å according to the golden-standard *Fourier* shell correlation (GSFSC) criterion. **b**. Angular distribution of the particles contributing to the final 3D reconstruction. The height of each spike indicated the number of particles in each designated orientation. **c**. Sharpened map of the $Ca_V2.3$ complex, colored according to the estimated value of local resolution. **d**. *Fourier* shell correlations (FSC) curves of the $Ca_V2.3$ complex and its atomic model. The unmasked (blue) and masked (red) FSC of the $Ca_V2.3$ EM map was calculated between two independently refined half-maps before and after post-processing. The model-vs-map FSC (black) was calculated between the full map and the atomic model. **e**. Representative cryo-EM density map (transparent grey surface) of the Domain II. Residues on the $S4_{II}$, $S6_{II}^{NCD}$ and W-helix were labeled. The lipid molecule residing near the $S4\text{-}S5_{II}$ was colored in green and also indicated.

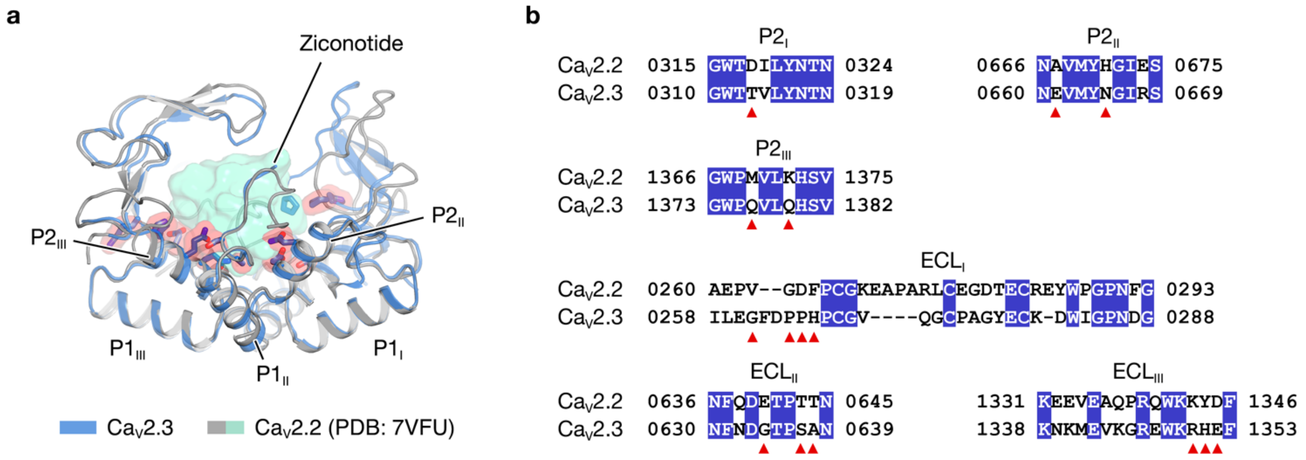


Supplementary Figure 3. Structural analysis of the ziconotide resistance of $Ca_V2.3$

**a**. Putative structural mismatches between the $Ca_V2.3$ and ziconotide. Extracellular helices (ECLs) and the P-loops (P1 and P2) of the $Ca_V2.3$ (blue) are superimposed on the corresponding structures of the ziconotide-bound $Ca_V2.2$. The ziconotide is shown as transparent green surface. Non-conserved residues that may contribute to the structural mismatch between $Ca_V2.3$ and ziconotide are shown as sticks, overlaid by red surfaces, and indicated in (**b**). **b**. Sequences alignments between $Ca_V2.3$ and $Ca_V2.2$ at the P-loops and ECLs. Conserved residues are highlighted in blue. Non-conserved residues that form close contact with ziconotide in $Ca_V2.2$ are indicated using red triangles.

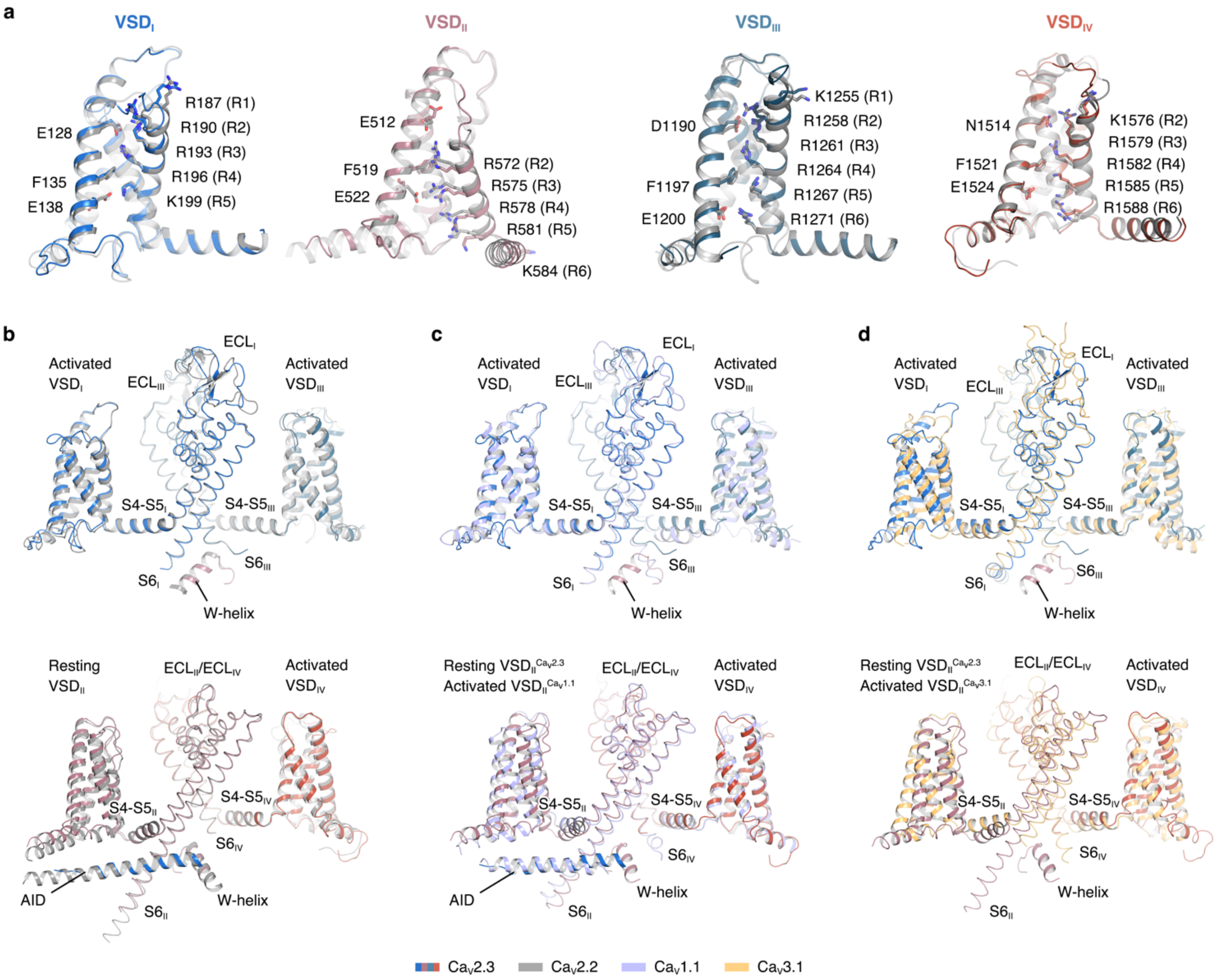


### Supplementary Figure 4. VSDs among the $Ca_V$ structures

Superimpositions of the voltage-sensing domains (VSDs) or pore domains between $Ca_V$2.3 and $Ca_V$2.2 (**a**–**b**), $Ca_V$1.1 (**c**) or $Ca_V$3.1 (**d**). The VSDs were shown as cartoon, and pore domain as loops. Four domains of $Ca_V$2.3 was colored blue, pink, deep cyan, and red, respectively. The $Ca_V$1.1, $Ca_V$2.2, and $Ca_V$3.1 was colored in light purple, gray, and yellow, respectively.

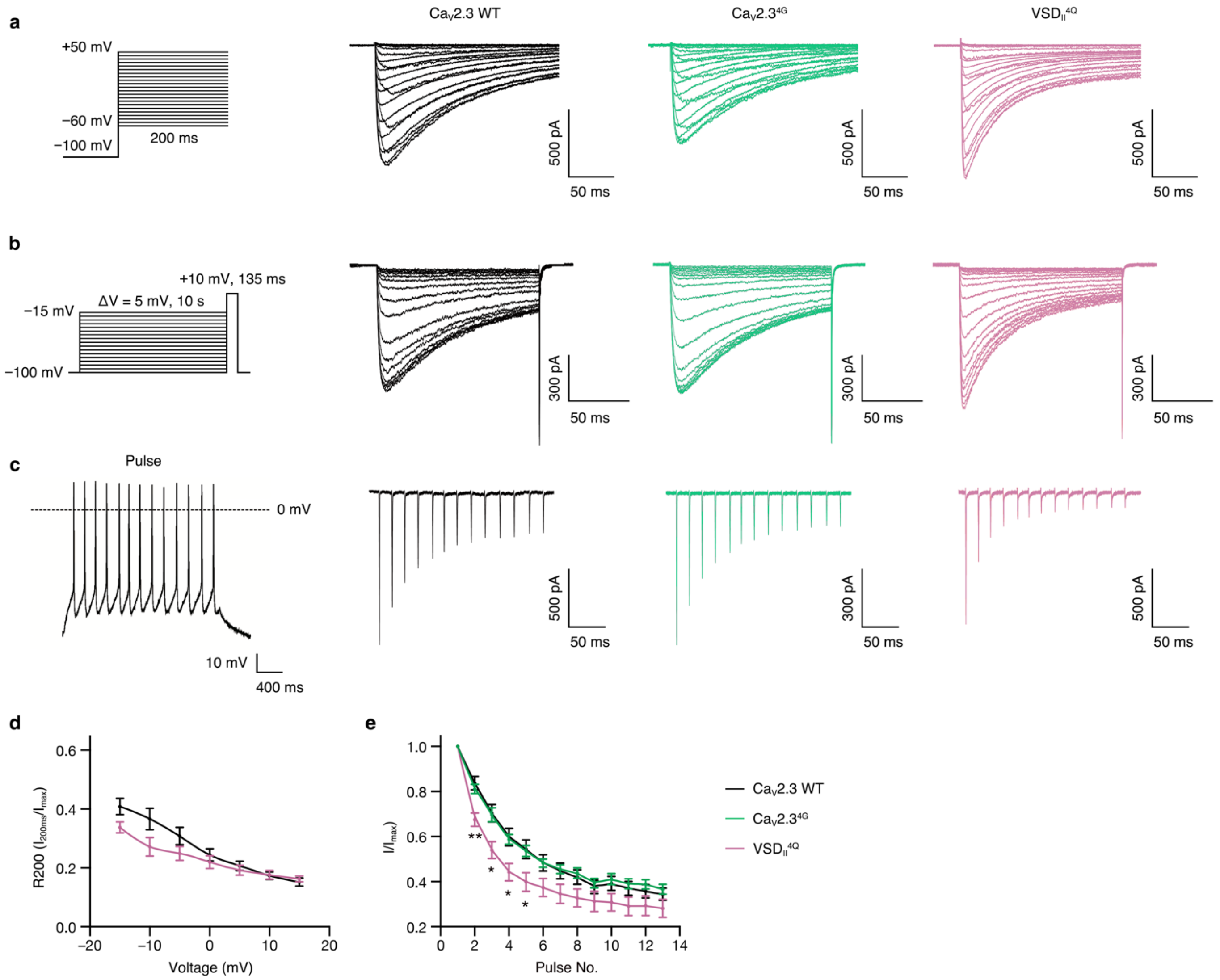


### Supplementary Figure 5. Representative whole-cell current traces for VSD-related mutants

Voltage-clamp protocols (left) and representative current traces for electrophysiological studies on the voltage-sensing domains (VSDs). **a**. Protocol and representative whole-cell voltage-clamp $Ca_V2.3$ currents (activation curves). Current traces were obtained from a series of 200-ms voltage steps from –60 mV to +50 mV, in 5-mV increments. Ratio of open-state inactivation (R200) were measured using 200-ms test pulses at +10 mV. **b**. Protocols and representative currents for inactivation curves. Currents were elicited by a +10-mV test pulse after holding-voltages from –100 mV to –15 mV, in 5-mV increments. **c**. Representative current responses stimulated by action potential (AP) trains (left). The AP trains were recorded using a whole-cell current-clamp from a mouse hippocampal CA1 pyramidal neuron after current injection (see Method section for the literature reference). **d**. R200 analysis of the mutants. $Ca_V2.3$ WT (black), n = 10; $VSD_{II}^{4Q}$ (pink), n = 7. **e**. Inactivation ratio of the mutants quantified using the current density (I) elicited by each spike of AP trains divided by the maximum current ($I_{max}$) elicited by the first spike. $Ca_V2.3$ WT (black), n = 13; $Ca_V2.3^{4G}$ (green), n = 7; $VSD_{II}^{4Q}$ (pink), n = 8. Significances were determined using two-sided, unpaired *t*-test. P values, $Ca_V2.3$ WT vs. $VSD_{II}^{4Q}$; 0.002 (No. 2), 0.01 (No. 3), 0.02 (No. 4), and 0.03 (No. 5).

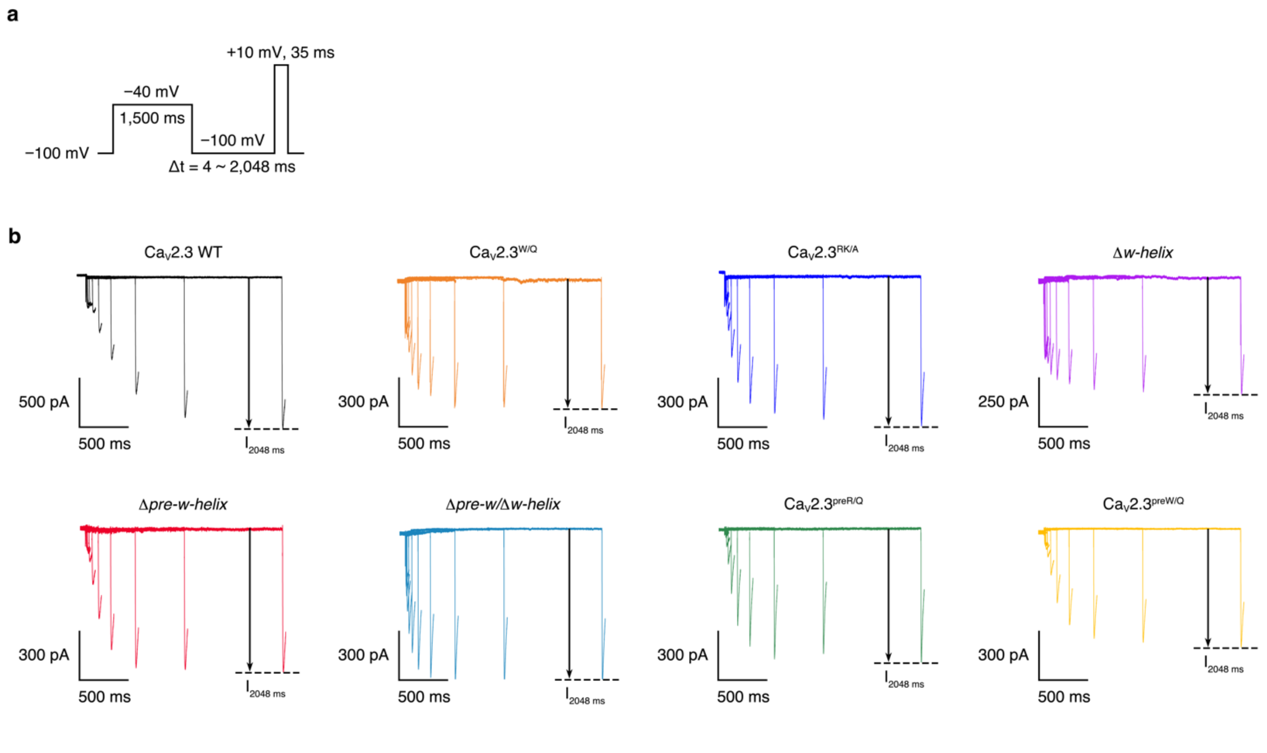


### Supplementary Figure 6. Representative whole-cell current traces for CSI-related mutants

Voltage-clamp protocol (**a**) and representative current traces (**b**) of the electrophysiological studies on the recovery rate from closed-state inactivation (CSI). HEK 293-T cells expressing $Ca_V2.3$ complex were depolarized using a –40 mV pre-pulse for 1,500 ms to inactivate the channels. A recovery hyperpolarization steps to –100 mV were subsequently applied for the indicated period (4–2,048 ms), followed by a 35-ms test pulse at +10 mV.

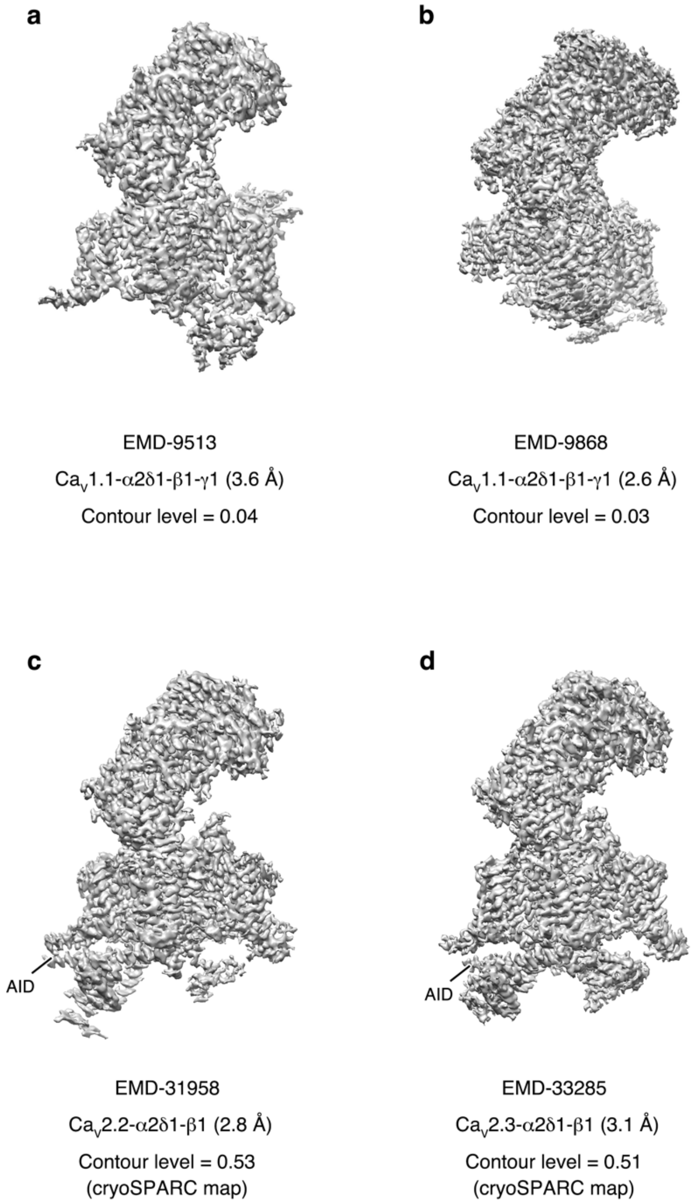


Supplementary Figure 7. Conformation heterogeneity of the AID

The alpha-interacting domains (AIDs) adopts a highly-dynamic conformation in the $Ca_V1.1$ structures (**a**–**b**) while were stabilized near the membrane plane in the intracellular side of the $Ca_V2.2$ structure (**c**) and the $Ca_V2.3$ structure (**d**) resolved in this study. The EM maps were shown as grey surfaces. Densities representing AIDs were labeled.

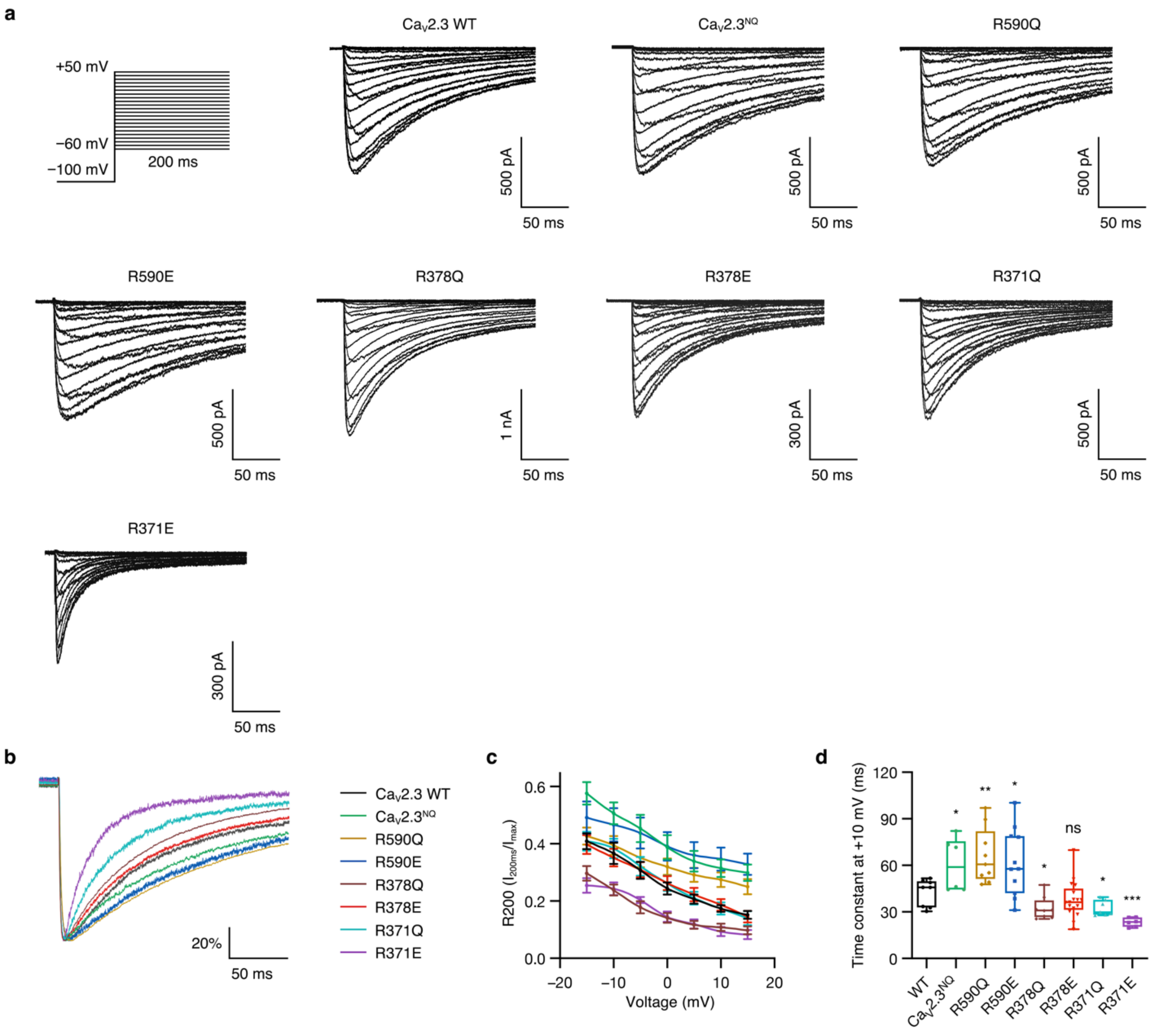


Supplementary Figure 8. Representative whole-cell current traces for OSI-related mutants

**a**. Voltage-clamp protocols and representative whole-cell current traces for electrophysiological studies on open-state inactivation (OSI). Current traces were obtained from a series of 200-ms voltage steps from –60 mV to +50 mV, in 5-mV increments. **b**. Representative whole-cell current traces of the OSI-related mutants under 200-ms test pulses at +10 mV. **c**. Ratio of OSI (R200) under the test pulse series ranging from –15 mV to 15 mV, measured using the current at the end of the 200-ms test pulse divided by the peak amplitude. $Ca_V2.3$ WT, n = 10; $Ca_V2.3^{NQ}$, n = 10; R590Q, n = 10; R590E, n = 6; R378Q, n = 9; R378E, n = 10; R371Q, n = 9; R371E, n = 7. **d**. Single-term exponential fitting of the current traces under 200-ms test pulses at +10 mV. $Ca_V2.3$ WT, n = 8; $Ca_V2.3^{NQ}$, n = 6; R590Q, n = 9; R590E, n = 11; R378Q, n = 8; R378E, n = 17; R371Q, n = 7; R371E, n = 6. Significances were determined using two-sided, unpaired *t*-test. P values, $Ca_V2.3$ WT vs. mutants; 0.02 ($Ca_V2.3^{NQ}$), 0.003 (R590Q), 0.03 (R590E), 0.03 (R378Q), 0.02 (R371Q), and 0.0003 (R371E).

**Supplementary Table 1.**

**Cryo-EM data collection, refinement, and validation statistics**

| | Cav2.3-α2δ1-β1 (EMD-33285) (PDB 7XLQ) |
|---|---|
| **Data collection and processing** | |
| Magnification | ×130,000 |
| Voltage (kV) | 300 |
| Electron exposure ($e^{–}/Å^2$) | 60 |
| Defocus range (μm) | −1.2 – −2.2 |
| Pixel size (Å) | 1.04 |
| Symmetry imposed | C1 |
| Initial particle images (no.) | 787,518 |
| Final particle images (no.) | 257,473 |
| Map resolution (Å) | 3.1 |
| FSC threshold | 0.143 |
| Map resolution range (Å) | 2.5 – 4.5 |
| **Refinement** | |
| Initial model used (PDB code) | 7VFS |
| Model resolution (Å) | 3.3 |
| FSC threshold | 0.5 |
| Map sharpening *B* factor ($Å^2$) | −94.7 |
| Model composition | |
| Non-hydrogen atoms | 20,247 |
| Protein residues | 2,436 |
| Ligands | 36 |
| *B* factors ($Å^2$) | |
| Protein | 67.32 |
| Ligand | 53.85 |
| R.m.s. deviations | |
| Bond lengths (Å) | 0.006 |
| Bond angles (°) | 0.805 |
| Validation | |
| MolProbity score | 2.03 |
| Clashscore | 10.84 |
| Poor rotamers (%) | 0.23 |
| Ramachandran plot | |
| Favored (%) | 92.38 |
| Allowed (%) | 7.58 |
| Disallowed (%) | 0.04 |
